# Pdgfab/Pdgfra-mediated chemoattraction guides the migration of sclerotome-derived fibroblast precursors in zebrafish

**DOI:** 10.1101/2024.09.24.614639

**Authors:** Emilio E. Méndez-Olivos, Katrinka M. Kocha, Shan Liao, Peng Huang

## Abstract

In vertebrates, the sclerotome is a transient embryonic structure that gives rise to various tissue support cells, including fibroblasts. However, how fibroblast precursors are guided to diverse tissues remain poorly understood. Using zebrafish, our lab has previously shown that sclerotome-derived cells undergo extensive migration to generate distinct fibroblasts subtypes, including tenocytes along the myotendinous junction and fin mesenchymal cells in the fin fold. Interestingly, the pan-fibroblast gene platelet-derived growth factor receptor a (*pdgfra*), which has been implicated in cell migration across various contexts, is specifically expressed in the sclerotome and its descendants. Loss of functional Pdgfra in a *pdgfra* gene-trap mutant results in severe defects in the migration of sclerotome-derived cells, leading to a dose-dependent loss of tenocytes and fin mesenchymal cells. By combining live imaging and mosaic labeling with a membrane-bound dominant-negative tool, we demonstrate that Pdgfra acts cell-autonomously to regulate the migration of sclerotome-derived cells. In the absence of ligand *pdgfab*, which is expressed in the medial somite, sclerotome-derived cells fail to migrate medially, resulting in a loss of tenocytes, although they can migrate normally toward the fin fold and generate fin mesenchymal cells. Strikingly, localized expression of Pdgfab in *pdgfab* mutants can direct the migration of sclerotome-derived cells to both normal and ectopic locations, suggesting a chemoattractive role for the Pdgfab ligand. Together, our results demonstrate that Pdgfab/Pdgfra-mediated chemoattraction guides the migration of sclerotome-derived fibroblast precursors to specific locations, where they diversify into distinct fibroblast subtypes.

## INTRODUCTION

Almost every organ in our body contains connective tissue, with fibroblasts being the main cellular component. Known primarily for producing the extracellular matrix, fibroblasts also perform various other functions, including secreting growth factors, creating niches for stem cells, modulating immune response, and contributing to wound healing (Younesi et al., 2024). The involvement of fibroblasts in such a wide range of processes raises the question of whether they are a single cell type. Thanks to single-cell sequencing technologies, many groups have reported that fibroblasts are substantially heterogeneous in many organs, including the heart, skin, skeletal muscle, intestine, bladder, and synovial joints (Collins et al., 2023; Driskell & Watt, 2015; Muhl et al., 2020; Tallquist, 2020). Although fibroblasts arise from all three germ layers during embryonic development (Plikus et al., 2021), little is known about the molecular mechanisms controlling how fibroblast precursors migrate to distinct locations and differentiate into diverse fibroblast subtypes. The somite is a transient embryonic structure that gives rise to the trunk of vertebrates.

In amniotes, the sclerotome is a sub-compartment formed in the ventral part of the somite (Draga & Scaal, 2024). Its development depends on signaling molecules derived from the notochord, with Sonic Hedgehog (SHH) being a key factor for its induction (Johnson et al., 1994). The sclerotome compartment later gives rise to the axial skeleton, including the vertebral body, ribs, and neural arches, as well as their associated tendons and ligaments (Scaal, 2016). Our previous studies show that the zebrafish sclerotome has a unique bipartite organization in the ventral and dorsal parts of the somite, specifically marked by the expression of *nkx3.1* (Ma et al., 2018). Live imaging, lineage tracing, and single-cell RNA sequencing experiments demonstrate that sclerotome progenitors undergo extensive migration to generate diverse fibroblast subtypes, including tenocytes, perivascular fibroblasts, and fin mesenchymal cells (Ma et al., 2018; Rajan et al., 2020, 2023).

Interestingly, although dispensable for sclerotome induction, active Shh signaling is required for the migration of sclerotome-derived cells toward the notochord (Ma et al., 2018). However, the exact mechanisms controlling this migration remain unexplored.

One molecular pathway often associated with fibroblasts is the platelet-derived growth factor (PDGF) signaling pathway. This pathway comprises four secreted ligands (PDGFA, PDGFB, PDGFC, and PDGFD) and two tyrosine kinase receptors (PDGFRA and PDGFRB) (Andrae et al., 2008; P.-H. Chen et al., 2013). PDGFRA is known as a pan-fibroblast marker (Horikawa et al., 2015) and controls the migration of various cells during embryonic development, such as cranial and cardiac neural crest cells in mouse and zebrafish (Eberhart et al., 2008; Tallquist & Soriano, 2003), myocardial precursors in zebrafish (Bloomekatz et al., 2017; El-Rass et al., 2017), and mesendodermal cells in Xenopus and zebrafish (Damm & Winklbauer, 2011; Montero et al., 2003). Interestingly, all these migratory events are mediated by the ligand PDGFA, which is thought to function as a chemoattractant for the PDGFRA-expressing cells, although this has been inferred from its expression pattern.

Homozygous *Pdgfra* knock-out mice die at E14-E16 with a cleft palate phenotype, consistent with the role of *Pdgfra* in neural crest cell migration (Soriano, 1997). These *Pdgfra* mutants also display various defects in vertebral neural arches, which are derived from the sclerotome (Pickett et al., 2008; Soriano, 1997). In contrast, homozygous *Pdgfa* knock-out mice are viable but die by week 6 postnatally, likely due to respiratory issues (Boström et al., 1996).

Interestingly, *Pdgfa* mutants display early somite patterning defects similar to those observed in *Pdgfra* null mice and *Pdgfa* somite expression promotes rib and vertebral development (Tallquist et al., 2000). These findings suggest that the PDGFA/PDGFRA signaling pathway plays a role in the development of the sclerotome lineage.

In this study, we describe the critical role of Pdgfra signaling in guiding the migration of sclerotome-derived fibroblast precursors. By combining mutant analysis with in vivo imaging, we demonstrate that *pdgfra* is required cell-autonomously for the migration of sclerotome-derived cells to different locations, where they differentiate into various fibroblast subtypes.

Interestingly, the ligand *pdgfab* is expressed in the medial somites in response to Shh signaling. Loss of *pdgfab* specifically impairs the migration of sclerotome-derived cells toward the notochord. Mosaic expression analysis further reveals that the Pdgfab ligand functions as a chemoattractant, guiding the migration of *pdgfra*-expressing cells. Our study suggests that region-specific expression of the Pdgfab ligand provides a localized cue to direct the migration of Pdgfra-expressing fibroblast precursors, thereby facilitating fibroblast diversification.

## RESULTS

### Characterization of a *pdgfra* gene-trap mutant

We previously showed that the zebrafish sclerotome has a bipartite organization occupying both the ventral and dorsal region of each somite along the trunk (Ma et al., 2018). Cells derived from both sclerotome domains undergo extensive migrations to give arise to different fibroblast populations (Ma et al., 2018, 2023). Interestingly, double fluorescent in situ hybridization (FISH) revealed that *pdgfra*, encoding platelet-derived growth factor receptor a, was co-expressed with sclerotome maker *nkx3.1* in both dorsal and ventral sclerotome domains, as well as sclerotome-derived cells around the notochord at 24 hours post-fertilization (hpf) (Fig. 1A). This result is consistent with our previous finding that *pdgfra* is broadly expressed across all fibroblast subtypes derived from the sclerotome (Rajan et al., 2023). Since *pdgfra* has been shown to regulate the migration of cardiomyocytes (Bloomekatz et al., 2017) and cranial neural crest cells (Eberhart et al., 2008) in zebrafish, we asked whether *pdgfra* is also required for the migration of sclerotome-derived cells. To test this, we characterized a *pdgfra* gene-trap line (El-Rass et al., 2017) with an insertion in the intron upstream of exon 17, which encodes the transmembrane domain (Fig. 1B). This gene-trap line is predicted to produce a non-functional truncated Pdgfra protein, lacking the transmembrane and kinase domains, fused with mRFP (PdgfraτιTK-mRFP) (Fig. 1B). We predicted that PdgfraτιTK-mRFP can bind to Pdgfra ligands in the extracellular space but is unable to signal due to the absence of transmembrane and kinase domains, thereby functioning as a dominant-negative decoy receptor. The gene-trap line is also predicted to generate ubiquitously expressed GFP fused with the Pdgfra transmembrane and kinase domains under the control of the *β-actin* promoter (Fig. 1B). For simplicity, we designate wild-type, heterozygous, and homozygous *pdgfra* fish as *pdgfra^+/+^*, *pdgfra^mRFP/+^*, and *pdgfra^mRFP/mRFP^*. Using region-specific probes, we confirmed that *mRFP* showed an identical pattern as the *pdgfra5’* probe in *pdgfra^mRFP/mRFP^* fish, while staining with *GFP* and *pdgfra3’* probes resulted in similar ubiquitous patterns in homozygous fish (Fig. S1).

**Figure 1:**
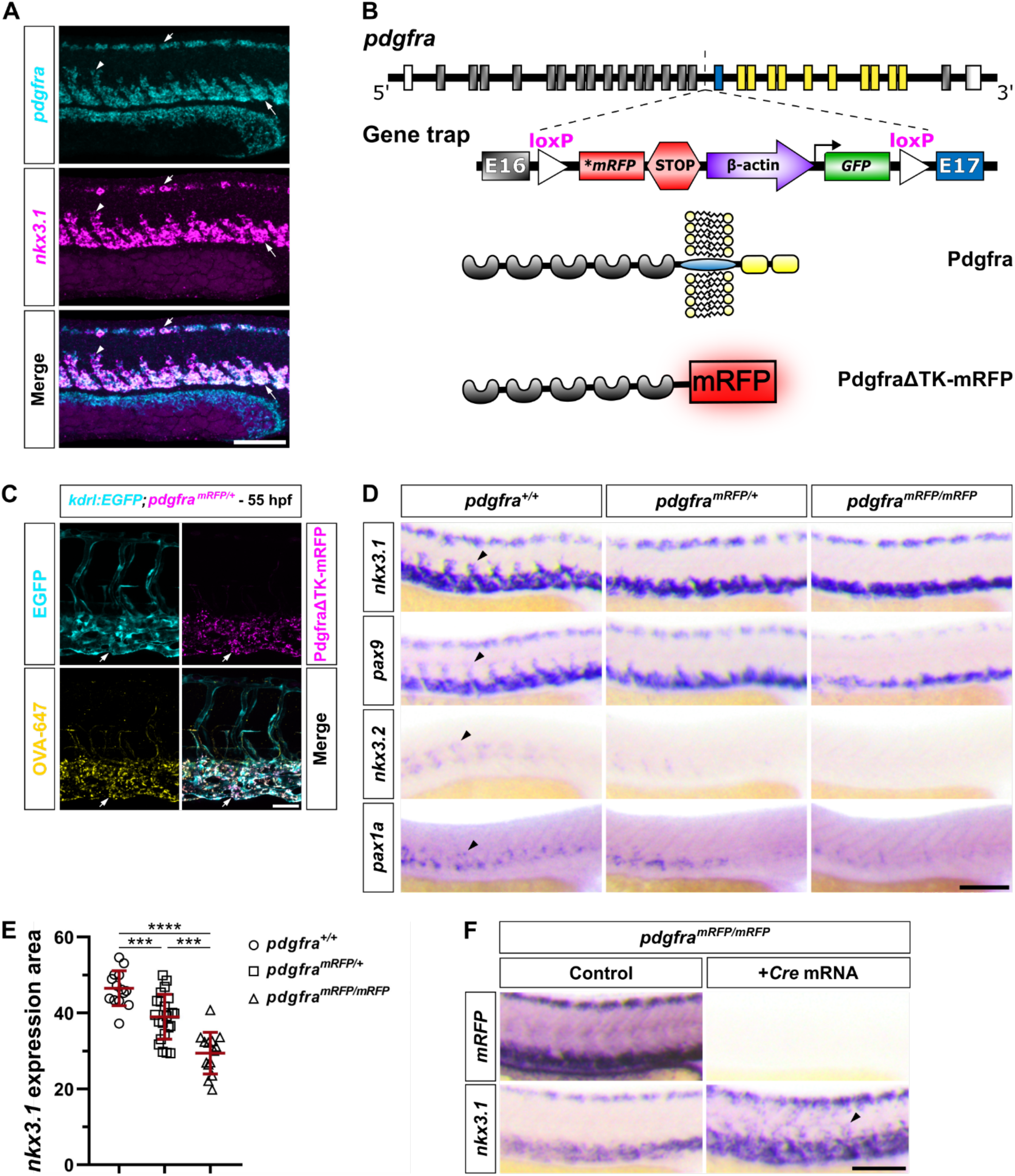
***pdgfra* is required for the migration of sclerotome-derived cells.** (A) Wild-type embryos at 24 hpf were co-labeled with *pdgfra* (cyan) and *nkx3.1* (magenta). *pdgfra* and *nkx3.1* are co-expressed in the dorsal sclerotome (short arrows), ventral sclerotome (long arrows), and sclerotome-derived notochord-associated cells (arrowheads). *n* = 12 embryos. (B) Schematics of the *pdgfra* gene-trap line. The exons of the *pdgfra* gene are color-coded: gray for extracellular coding exons, blue for the transmembrane coding exon, yellow for intracellular coding exons, and white for non-coding exons. The gene-trap insertion is flanked by two loxP sites and is located in the intron between exons 16 and 17. The gene-trap mutant is predicted to produce a secreted protein containing the extracellular domain of Pdgfra fused with mRFP (PdgfraΔTK-mRFP). (C) *kdrl:EGFP; pdgfra^mRFP/+^* embryos were injected with Alexa Fluor 647-conjugated ovalbumin (OVA-647) into the extracellular space at the blastula stage (4 hpf) and imaged at 55 hpf. Co-localization of OVA-647 (yellow) and PdgfraΔTK- mRFP (magenta) proteins in endothelial cells of the caudal vein plexus, labeled by the *kdrl:EGFP* reporter (cyan), are indicated by arrows. *n* = 20 embryos. (D) Expression analysis of sclerotome markers (*nkx3.1* and *pax9*) and sclerotome-derived notochord-associated cell markers (*nkx3.2* and *pax1a*) in *pdgfra* mutants at 24 hpf. The expression of all markers shows a dose-dependent reduction in the migrating sclerotome-derived cells (arrowheads) around the notochord across different mutant backgrounds. *n* = 25 embryos per staining. (E) Quantification of *nkx3.1* expression area in the experiments showed in (D). *n* = 14 (*pdgfra^+/+^*), 23 (*pdgfra^mRFP/+^*), and 14 (*pdgfra^mRFP/mRFP^*) embryos. Data are plotted as mean ± SD. Statistics: One-way ANOVA with Tukey’s multiple comparison test. Asterisks representation: *p* < 0.001 (***) and *p* < 0.0001 (****). (F) *pdgfra^mRFP/mRFP^*embryos injected with *Cre* mRNA at the one-cell stage, along with their uninjected controls, were stained for *mRFP* and *nkx3.1* expression at 24 hpf. *Cre*-expressing *pdgfra^mRFP/mRFP^* embryos show a complete loss of *mRFP* expression and restoration of migration of *nkx3.1^+^*sclerotome-derived cells (arrowhead) at 24 hpf. *n* = 25 embryos per staining. Scale bars: (A, D, F) 100 μm; (C) 50 μm.

We next performed live imaging to analyze the distribution of different fusion proteins generated from the gene-trap line. While the GFP fusion protein was undetectable, we found that PdgfraτιTK-mRFP puncta were specifically enriched in endothelial cells of the caudal vein plexus (CVP), labeled by *kdrl:EGFP* (Fig. 1C). Since endothelial cells do not normally express *pdgfra* and PdgfraτιTK-mRFP is predicted to be secreted, we hypothesized that scavenger endothelial cells (Campbell et al., 2018) phagocytose the secreted PdgfraτιTK-mRFP fusion protein from the extracellular space. To test this possibility, we injected Alexa Fluor 647- conjugated ovalbumin (OVA-647) into the extracellular space of *pdgfra^mRFP/+^; kdrl:EGFP* embryos at 4 hpf. Interestingly, both PdgfraτιTK-mRFP and OVA-647 proteins displayed a similar puncta pattern in CVP endothelial cells at 55 hpf (Fig. 1C). Together, our results suggest that the *pdgfra* gene-trap line produces a secreted PdgfraτιTK-mRFP protein, which is cleared from the extracellular space by scavenger endothelial cells.

### *pdgfra* mutants show defects in the migration of sclerotome-derived cells

To determine of the role of *pdgfra* in the migration of sclerotome-derived cells, we analyzed the expression pattern of sclerotome markers, *nkx3.1* and *pax9*, in *pdgfra* mutants at 24 hpf. Although both markers were normally expressed in the dorsal and ventral sclerotome domains, the expression of *nkx3.1* and *pax9* in sclerotome-derived notochord-associated cells was completely lost in *pdgfra^mRFP/mRFP^*fish (Fig. 1D). *pdgfra^mRFP/+^* fish also displayed a substantial reduction of *nkx3.1* and *pax9* staining around the notochord (Fig. 1D), consistent with the prediction that PdgfraτιTK-mRFP likely functions as a dominant decoy receptor. Indeed, quantification of *nkx3.1* expression area in *pdgfra^+/+^*, *pdgfra^mRFP/+^*, and *pdgfra^mRFP/mRFP^* embryos showed a significant reduction in a dose-dependent manner (Fig. 1E). The sclerotome-derived cells around the notochord can be specifically labeled by the expression of *nkx3.2* and *pax1a* (Ma et al., 2018). We observed a similar dose-dependent reduction in the expression of both markers in the *pdgfra* mutant background (Fig. 1E). To further validate these results, we removed the loxP-flanked gene-trap insertion (Fig. 1B) by injecting *Cre* mRNA into *pdgfra^mRFP/mRFP^* embryos at the one-cell stage. Early *Cre* expression resulted in the complete loss of *mRFP* expression in *pdgfra^mRFP/mRFP^* fish (Fig. 1F), suggesting that both copies of the insertion were successfully excised. Strikingly, the removal of the gene-trap insertion led to a substantial rescue of *nkx3.1*^+^ sclerotome-derived cells around the notochord in *pdgfra^mRFP/mRFP^*fish (Fig. 1F). Together, our results suggest that sclerotome-derived cells require functional Pdgfra to migrate toward the notochord region.

### Failed migration of sclerotome-derived cells impairs fibroblast development

The absence of sclerotome marker expression around the notochord suggests that cells fail to migrate out of the sclerotome domains. To directly visualize the migration of sclerotome-derived cells, we performed live imaging of *pdgfra* mutants from 22 to 36 hpf using the sclerotome reporter line, *nkx3.1:Gal4; UAS:NTR-mCherry* (Ma et al., 2018). In *pdgfra^+/+^* controls, mCherry^+^ cells emerged from both the dorsal and ventral sclerotome domains and migrated extensively, occupying the medial region of the entire trunk (Fig. 2A and Movie S1). By contrast, both *pdgfra^mRFP/+^* and *pdgfra^mRFP/mRFP^*embryos displayed severely reduced migration of mCherry^+^ cells (Fig. 2A and Movie S1). These results suggest that *pdgfra* is required for the migration of sclerotome-derived cells.

**Figure 2:**
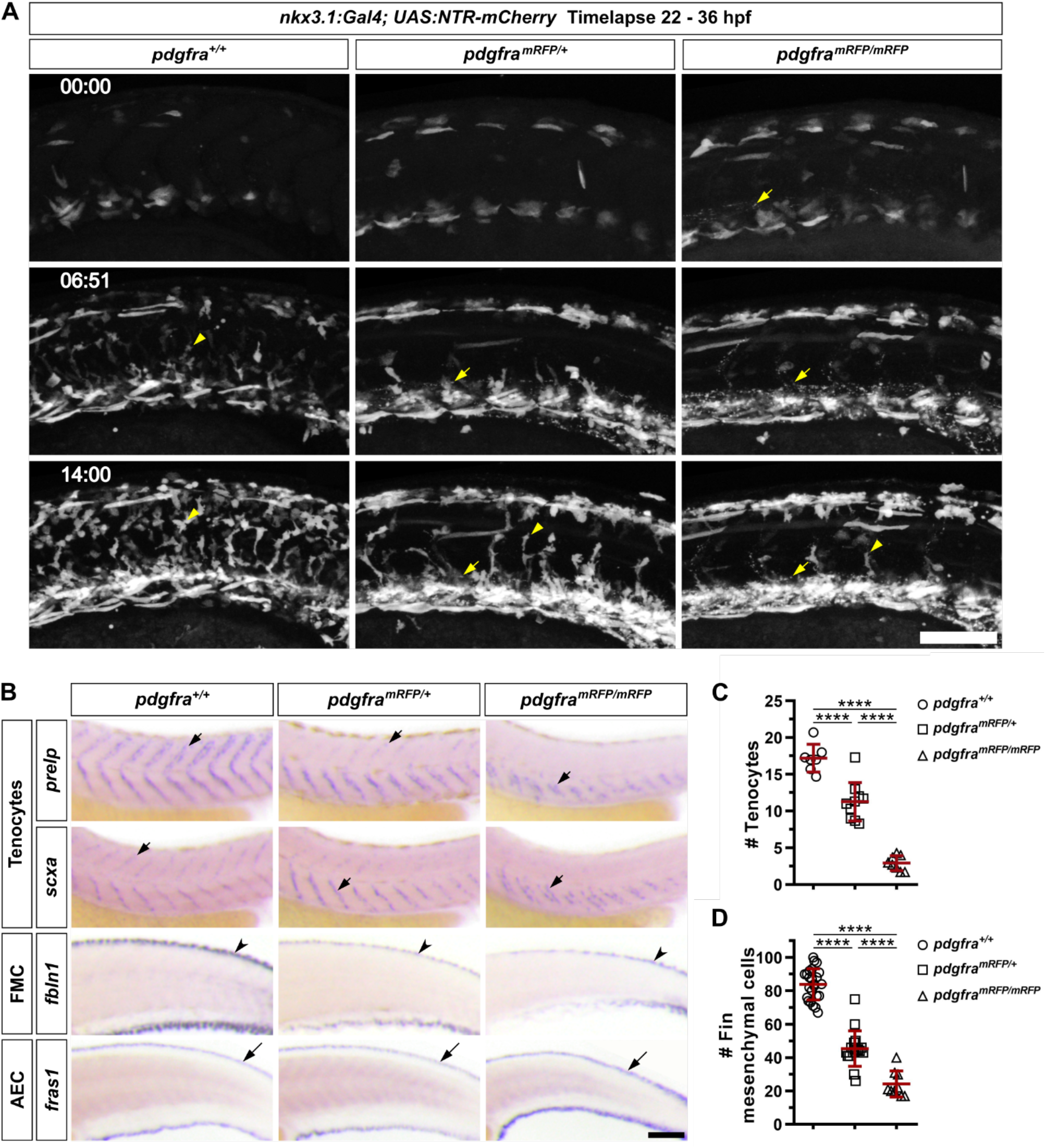
*pdgfra* mutants show defects in the generation of sclerotome-derived cells. (A) Snapshots from time-lapse imaging of the *nkx3.1:Gal4; UAS:NTR-mCherry* reporter in *pdgfra^+/+^*, *pdgfra^mRFP/+^*, and *pdgfra^mRFP/mRFP^* backgrounds from 22 to 36 hpf. While mCherry^+^ cells (arrowheads) migrate normally in wild-type siblings, both heterozygous and homozygous *pdgfra* mutants display substantially reduced migration, with *pdgfra^mRFP/mRFP^*mutants being the most affected. Note that the PdgfraΔTK-mRFP fusion proteins appear as bright dots (arrows) within the vasculature of *pdgfra^mRFP/+^*and *pdgfra^mRFP/mRFP^* embryos. The corresponding movies are shown in Movie S1. Time stamps are indicated in the hh:mm format. *n* = 4 embryos per group. (B) Expression of tenocyte markers (*prelp* and *scxa*), fin mesenchymal cell (FMC) marker (*fbln1*), and apical epidermal cell (AEC) marker (*fras1*) in *pdgfra^+/+^*, *pdgfra^mRFP/+^*, and *pdgfra^mRFP/mRFP^* embryos at 48 hpf. The expression of both tenocyte (short arrows) and fin mesenchymal cell (notched arrowheads) markers is reduced in the *pdgfra* mutant background, while the apical epidermal cell marker (long arrows) remains unchanged. *n* = 25 embryos per staining. (C) Quantifications of tenocytes using the *nkx3.1:Gal4; UAS:NTR-mCherry* reporter in embryos with different *pdgfra* genotypes at 48 hpf. Each dot represents the average number of mCherry^+^ tenocytes along one MTJ, based on counts from three junctions of a single embryo. *n* = 7 (*pdgfra^+/+^*), 9 (*pdgfra^mRFP/+^*), and 8 (*pdgfra^mRFP/mRFP^*) embryos. (D) Quantifications of fin mesenchymal cells in the major fin fold of embryos with different *pdgfra* genotypes at 48 hpf. Each dot represents the total number of fin mesenchymal cells from both the dorsal and ventral fin folds in the 8-somite region posterior to the end of the yolk extension in one embryo. *n* = 24 (*pdgfra^+/+^*), 17 (*pdgfra^mRFP/+^*), and 9 (*pdgfra^mRFP/mRFP^*) embryos. All data are plotted as mean ± SD. Statistics: One-way ANOVA with Tukey’s multiple comparison test. Asterisks representation: *p* < 0.0001 (****). Scale bars: 100 μm.

Based on our previous work (Ma et al., 2023), we predicted that the sclerotome progenitors need to migrate out of the sclerotome domains to differentiate into distinct fibroblast subtypes. To test this, we stained *pdgfra* mutants at 48 hpf with fibroblast subtype- specific markers, including *prelp* and *scxa* for tenocytes and *fbln1* for fin mesenchymal cells. In *pdgfra^+/+^* controls, tenocytes expressing *prelp* and *scxa* were present along the entire V- shaped myotendinous junction (MTJ) (Fig. 2B). By contrast, *pdgfra^mRFP/+^*embryos almost completely lacked tenocytes in the dorsal half of the MTJ, while *pdgfra^mRFP/mRFP^* fish showed further reduction of tenocytes, with only some remaining in the very ventral region of the MTJ (Fig. 2B). Quantification using *nkx3.1:Gal4; UAS:NTR-mCherry* showed a dose-dependent decrease in tenocyte number across different mutant backgrounds (Fig. 2C), reminiscent of our results with sclerotome markers (Fig. 1D-E). Similar to tenocytes, we also observed a dose-dependent loss of *fbln1^+^* fin mesenchymal cells in the fin folds (Fig. 2B and 2D). In contrast to *fbln1*, we observed no difference in the staining of *fras1*, a marker for apical epidermal cells that are not sclerotome-derived (Fig. 2B), suggesting that the fin fold compartment is formed normally. Together, our results suggest that the migration defect caused by the loss of *pdgfra* compromises the generation of sclerotome-derived fibroblasts, including tenocytes and fin mesenchymal cells.

### Pdgfra functions cell-autonomously in regulating the migration of sclerotome-derived cells

To validate the *pdgfra* mutant results and assess the autonomy of Pdgfra in the migration of sclerotome-derived cells, we generated two dominant-negative forms of the receptor. In the first construct, we used the heat shock promoter to drive the expression of the extracellular domain of Pdgfra fused with EGFP (*hsp:pdgfraτιTK-EGFP*), mimicking the secreted PdgfraτιTK-mRFP decoy receptor produced from the gene-trap line (Fig. 3A). As a control, we expressed secreted GFP under the heat shock promoter (*hsp:sec-GFP*) (Yamaguchi et al., 2019) (Fig. 3B). Induction of the secreted PdgfraτιTK-EGFP decoy receptor, but not the secreted EGFP, at 18 hpf, effectively abrogated the migration of *nkx3.1^+^* sclerotome-derived cells at 24 hpf (Fig. 3B-C). This result suggests that a functional Pdgfra receptor is required for the migration of sclerotome-derived cells, confirming our *pdgfra* mutant results.

**Figure 3:**
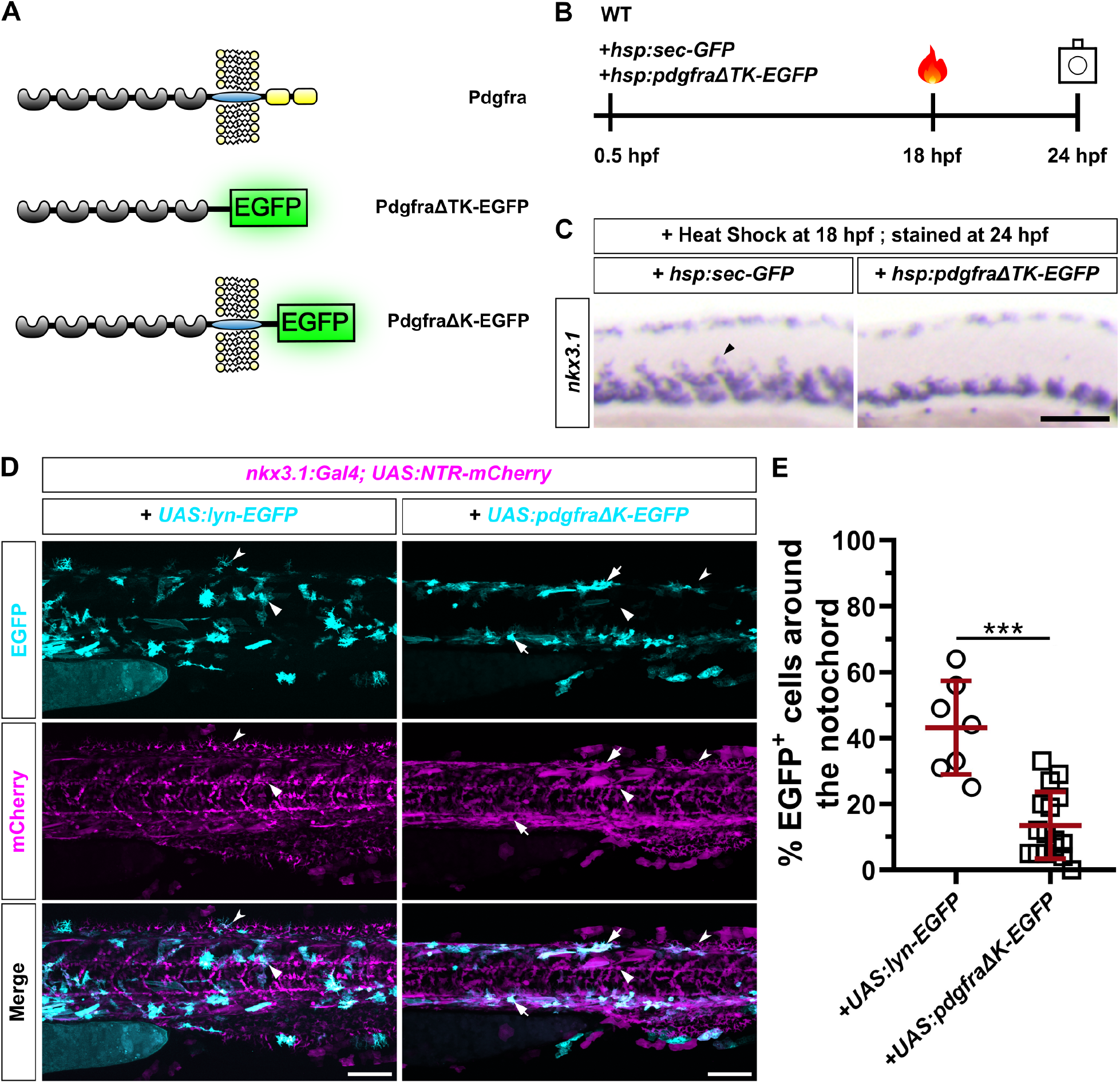
*pdgfra* is required cell-autonomously for sclerotome-derived cell migration. (A) Schematics of endogenous Pdgfra protein and two different loss-of-function variants. Compared to wild-type Pdgfra protein (top), PdgfraΔTK-EGFP (middle) is a secreted decoy receptor with its transmembrane and cytoplasmic domains replaced by EGFP, while PdgfraΔK-EGFP (bottom) is a membrane-bound dominant-negative receptor with only the cytoplasmic domain replaced by EGFP. (B) Schematics of overexpression experiments. Wild- type embryos were injected at the one-cell stage with either *hsp:sec-GFP* or *hsp:pdgfraΔTK- EGFP* plasmids, heat shocked at 18 hpf, and stained and imaged at 24 hpf. (C) Expression of *nkx3.1* in embryos at 24 hpf after overexpression of either secreted GFP or PdgfraΔTK- EGFP. Ectopic expression of PdgfraΔTK-EGFP, but not secreted GFP, impairs the migration of *nkx3.1*^+^ cells from the sclerotome (arrowhead). *n* = 20 embryos per staining. (D) Mosaic expression of Lyn-EGFP or PdgfraΔK-EGFP (cyan) in *nkx3.1:Gal4; UAS:NTR-mCherry* (magenta) embryos at 50 hpf. In Lyn-EGFP-expressing embryos, EGFP^+^ cells are widely distributed in the trunk both around the notochord (arrowheads) and in the fin fold (notched arrowheads). By contrast, in PdgfraΔK-EGFP-expressing embryos, while mCherry^+^EGFP^-^ cells can be found around the notochord (arrowheads) and in the fin fold (notched arrowheads), EGFP^+^ cells (arrows) are mostly restricted in the dorsal or ventral regions of the trunk. *n* = 10 embryos per group. (E) Quantification of the percentage of EGFP^+^ cells around the notochord from the experiments shown in (D). *n* = 7 (*UAS:lyn-EGFP*) and 16 (*UAS:pdgfraΔK-EGFP*) embryos. Data are plotted as mean ± SD. Statistics: Unpaired t test with Welch’s correction. Asterisks representation: *p* < 0.001 (***). Scale bars: 100 μm.

To determine whether Pdgfra acts cell-autonomously in sclerotome-derived cells, we developed a membrane-tethered dominant-negative Pdgfra receptor. We used the UAS driver to control the expression of a truncated Pdgfra with the kinase domain replaced by EGFP (*UAS:pdgfraτιK-EGFP*) (Fig. 3A). We mosaically expressed either *UAS:lyn-EGFP* (membrane-localized EGFP) or *UAS:pdgfraτιK-EGFP* in *nkx3.1:Gal4; UAS:NTR-mCherry* embryos. *UAS:lyn-EGFP*-injected fish showed a wide distribution of EGFP^+^ sclerotome- derived cells at 48 hpf, including cells around the notochord and fin mesenchymal cells in the fin fold (Fig. 3D). By contrast, in *UAS:pdgfraτιK-EGFP*-injected embryos, most of the mCherry^+^EGFP^+^ cells remained in either the ventral or dorsal region of the trunk, whereas mCherry^+^EGFP^-^ cells were distributed normally throughout the medial trunk (Fig. 3D).

Compared to *UAS:lyn-EGFP*, expression of *UAS:pdgfraτιK-EGFP* resulted in a significant decrease in the percentage of EGFP^+^ cells around the notochord (Fig. 3E). Together, these results suggest that Pdgfra acts cell-autonomously in controlling the migration of sclerotome- derived cells.

### Hedgehog signaling regulates the migration of sclerotome-derive cells by controlling ***pdgfab* expression**

Mammalian PDGFRA receptor is regulated by PDGF ligands, including PDGFA and PDGFC (Fredriksson et al., 2004). To identify the ligands that regulate the migration of sclerotome-derived cells, we performed in situ hybridization to examine the expression pattern of the three candidate ligands, *pdgfaa*, *pdgfab*, and *pdgfc*. In wild-type embryos at 24 hpf, *pdgfab* displayed the most notable pattern in the medial trunk, with expression in the medial somites, motoneurons, and the hypochord (Fig. 4A-D). Meanwhile, *pdgfaa* was predominantly expressed in the dorsal spinal cord and the hypochord, while *pdgfc* was strongly expressed along the edge of the fin fold and in some anterior somites (Fig. 4A). The expression of *pdgfab* in the medial region of the trunk suggests that Pdgfab might regulate the migration of sclerotome-derived cells to surround the notochord. Indeed, co-staining of *pdgfab* and *pdgfra* revealed a complementary pattern, where *pdgfra^+^* sclerotome-derived cells were sandwiched between *pdgfab^+^* somitic cells on one side and *pdgfab*-expressing motoneurons and hypochord on the other (Fig. 4B). Double labeling with *pdgfab* and *myod1*, a myotome- specific marker, further confirmed that the medial part of the *myod1^+^* myotome co-expressed *pdgfab* (Fig. 4C).

**Figure 4:**
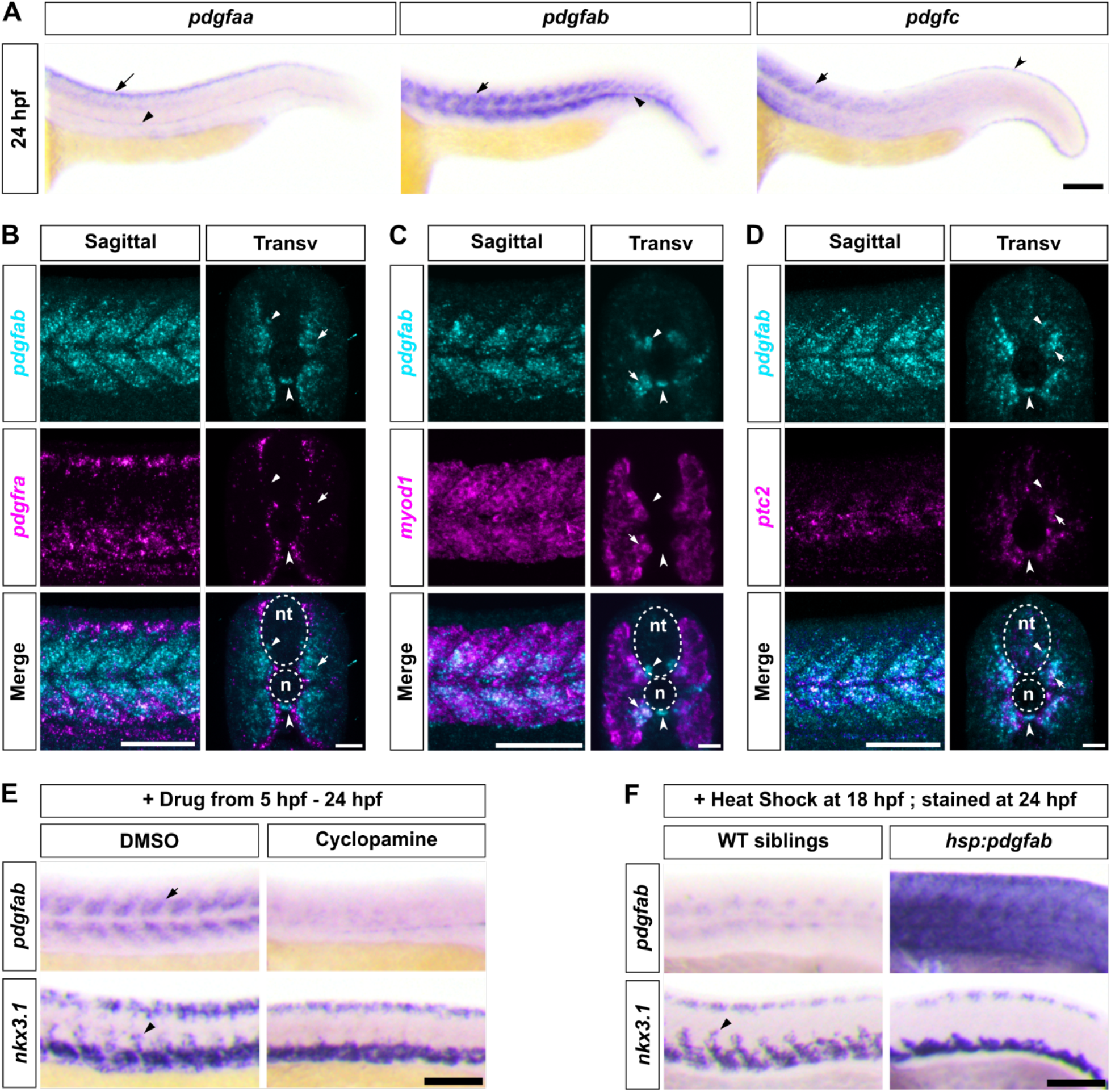
Hh signaling controls sclerotome-derived cell migration by regulating the expression of *pdgfab*. (A) Expression patterns of Pdgfra ligands in wild-type embryos at 24 hpf. *pdgfaa* expression is detected in the dorsal spinal cord (long arrow) and hypochord (arrowhead). *pdgfab* is expressed in the medial regions of the somites (arrow) and the hypochord (arrowhead). *pdgfc* expression is observed in anterior somites (arrow) and along the edge of the fin fold (notched arrowhead). *n* = 15 embryos per staining. (B-D) Wild-type embryos at 24 hpf were co-labeled with *pdgfab* (cyan) and *pdgfra* (B), *myod1* (C), or *ptc2* (D) (magenta). *pdgfab* is expressed in medial somites (arrows), motoneurons (arrowheads), and the hypocord (notched arrowhead). Sagittal and transverse views are shown with neural tube (nt) and notochord (n) indicated by dotted lines. *n* = 15 embryos per group. (B) *pdgfab* shows complementary expression to *pdgfra*. (C) *pdgfab* is expressed in the medial-most *myod1^+^* somitic region (arrows) adjacent to the notochord. (D) *pdgfab* and *ptc2* show co-expression in somitic cells (arrows) surrounding the notochord. (E) Expression analysis of *pdgfab* and *nkx3.1* in wild-type embryos treated with DMSO or cyclopamine from 5 to 24 hpf. Cyclopamine treatment leads to near-complete loss of *pdgfab* expression in somites (arrow) and absence of *nkx3.1^+^* sclerotome-derived cells (arrowhead) surrounding the notochord. *n* = 15 embryos per group. (F) *hsp:pdgfab-2A-mCherry* transgenic embryos and their wild-type siblings were heat shocked at 18 hpf and analyzed for *pdgfab* and *nkx3.1* expression at 24 hpf. Compared to wild-type controls, *hsp:pdgfab-2A-mCherry* embryos show strong and ubiquitous *pdgfab* expression and impaired migration of *nkx3.1*-expressing cells (arrowhead) toward the notochord. *n* = 15 embryos per group. Scale bars: (A, E, F) 100 μm; (B, C, D) 100 μm (sagittal) and 50 μm (transverse).

The *pdgfab* expression in the medial somite is reminiscent of the pattern of Hedgehog (Hh) signaling target genes, such as *ptc2* and *gli1* (Concordet et al., 1996; Karlstrom et al., 2003). Indeed, double labeling of *pdgfab* and *ptc2* revealed a partially overlapping expression pattern, particularly in somitic cells close to the notochord, although *pdgfab* had a slightly broader expression domain than *ptc2* (Fig. 4D). The expression analysis suggests that *pdgfab* expression may be regulated by Hh signaling. To test this possibility, we treated wild-type embryos with DMSO or cyclopamine, a specific inhibitor of Hh signaling, from 5 to 24 hpf and stained them with *pdgfab* and *nkx3.1*. Strikingly, cyclopamine-treated fish showed an almost complete loss of *pdgfab* expression in the medial trunk, accompanied by the loss of *nkx3.1^+^* sclerotome-derived cells around the notochord (Fig. 4E). These results suggest that Hh signaling regulates the migration of sclerotome-derive cells by controlling *pdgfab* expression. To determine how Pdgfab regulates the migration of sclerotome-derive cells, we overexpressed Pdgfab using a *hsp:pdgfab-2A-mCherry* transgenic line. Ubiquitous expression of *pdgfab* induced by heat shock at 18 hpf resulted in a complete loss of migrating *nkx3.1^+^*sclerotome-derived cells at 24 hpf (Fig. 4F). This result suggests that Pdgfab likely acts an instructive signal in controlling the migration of sclerotome derivatives.

### *pdgfab* mutants display defective migration of sclerotome-derived cells to the notochord region

If Pdgfab is responsible for the migration of *nkx3.1*^+^ sclerotome-derived cells, we predicted that *pdgfab* loss-of-function mutants should partially phenocopy our *pdgfra* mutants. We characterized a nonsense mutant, *pdgfab^sa11805^*, generated by the Zebrafish Mutation Project at the Wellcome Sanger Institute (Kettleborough et al., 2013). The *pdgfab^sa11805^*mutation introduces a premature stop codon, resulting in a truncated peptide of 111 amino acids (aa) instead of 230 aa. This shortened peptide is predicted to have the Pdgfab N- terminal domain, but it lacks the dimerization domain and most of the predicted receptor binding interface. For simplicity, we designate wild-type, heterozygous, and homozygous fish as *pdgfab^+/+^*, *pdgfab^+/-^*, and *pdgfab^-/-^*, respectively. Using the *nkx3.1:Gal4*; *UAS:NTR-mCherry* reporter, we performed time-lapse imaging of sclerotome-derived cells from 24 to 34 hpf in different *pdgfab* mutant backgrounds (Fig. 5A and Movie S2). By 30 hpf, mCherry^+^ cells had dispersed throughout the medial trunk in *pdgfab^+/+^*controls, whereas in *pdgfab^-/-^* embryos, only a few mCherry^+^ cells had begun to emerge from the sclerotome domains (Fig. 5A and Movie S2). At 34 hpf, *pdgfab^-/-^* embryos showed a considerably less coverage of mCherry^+^ cells around the notochord compared to *pdgfab^+/+^*or *pdgfab^+/-^* fish (Fig. 5A and Movie S2).

**Figure 5:**
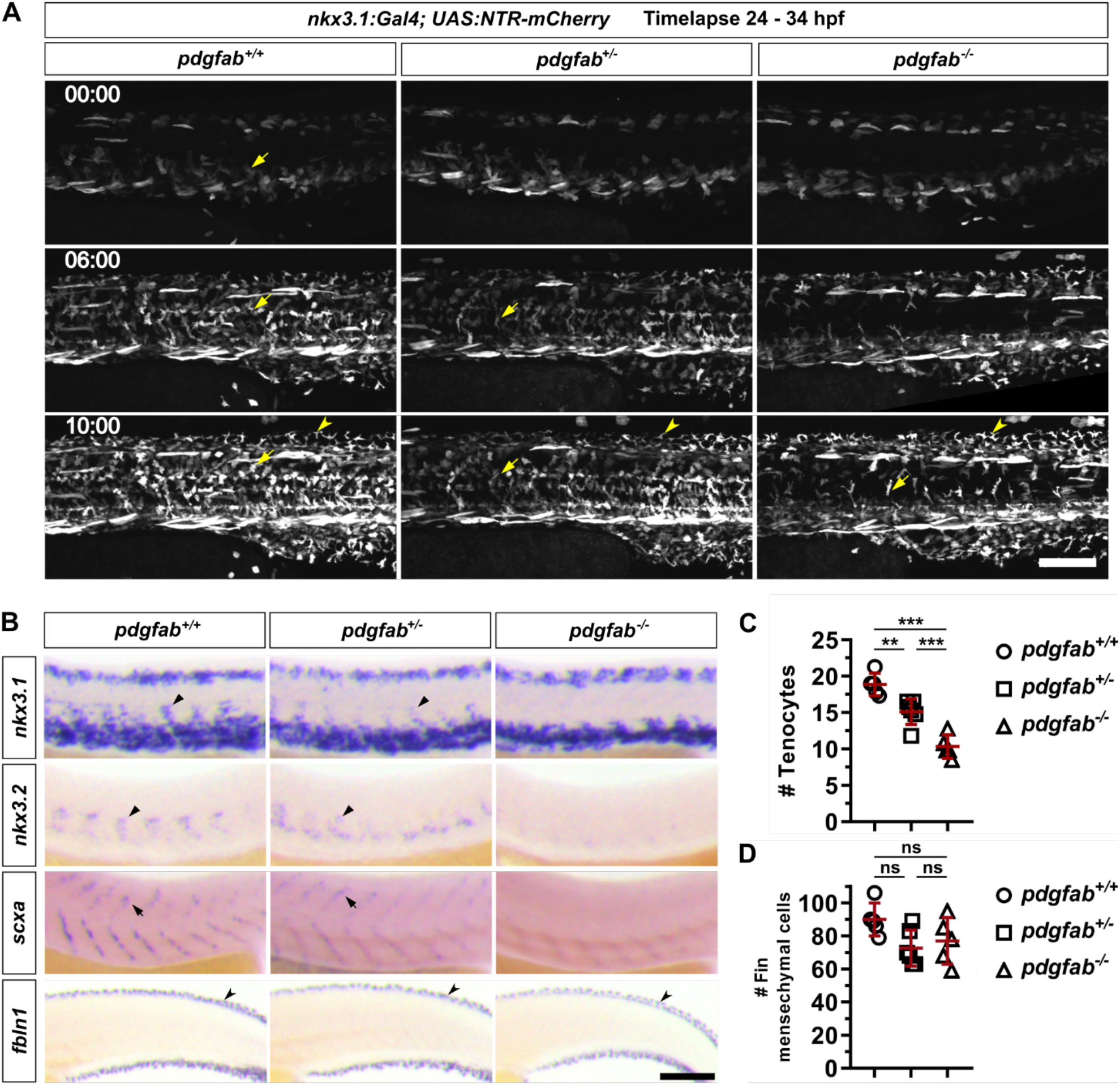
***pdgfab* is required for the migration of sclerotome-derived cells to the notochord.** (A) Snapshots from time-lapse imaging of the *nkx3.1:Gal4; UAS:NTR-mCherry* reporter in *pdgfab^+/+^*, *pdgfab^+/-^*, and *pdgfab^-/-^*backgrounds from 24 to 34 hpf. In *pdgfab^-/-^* embryos, the migration of mCherry^+^ cells (arrows) to the notochord is severely delayed compared to wild-type and heterozygous fish, while cells migrate normally to the fin fold to generate fin mesenchymal cells (notched arrowheads). The corresponding movies are shown in Movie S2. Time stamps are indicated in the hh:mm format. *n* = 4 embryos per group. (B) Marker analysis in embryos with different *pdgfab* genotypes. *pdgfab^-/-^*embryos lack *nkx3.1* and *nkx3.*2 expression in sclerotome-derived notochord-associated cells (arrowheads) at 24 hpf. At 48 hpf, the tenocyte marker *scxa* (arrows) is greatly reduced in *pdgfab^-/-^*embryos, while the fin mesenchymal cell marker *fbln1* (notched arrowheads) remains unchanged across all genotypes. *n* = 25 embryos per staining. (C) Quantifications of tenocytes using the *nkx3.1:Gal4; UAS:NTR-mCherry* reporter in embryos with different *pdgfab* genotypes at 56 hpf. Each dot represents the average number of mCherry^+^ tenocytes along one MTJ, based on counts from three junctions of an individual embryo. *n* = 5 (*pdgfab^+/+^*), 6 (*pdgfab^+/-^*), and 5 (*pdgfab^-/-^*) embryos. (D) Quantifications of fin mesenchymal cells in the major fin fold of embryos with different *pdgfab* genotypes at 56 hpf. Each dot represents the total number of fin mesenchymal cells from both the dorsal and ventral fin folds in the 8-somite region posterior to the end of the yolk extension in one embryo. *n* = 5 (*pdgfab^+/+^*), 6 (*pdgfab^+/-^*), and 5 (*pdgfab^-/-^*) embryos. Data are plotted as mean ± SD. Statistics: One-way ANOVA with Tukey’s multiple comparison test. Asterisks representation: *p* < 0.001 (***), *p* < 0.01 (**), and *p* > 0.05 (ns, not significant). Scale bars: 100 μm.

Interestingly, the migration of mCherry^+^ sclerotome-derived cells toward the fin fold seemed unaffected in *pdgfab^-/-^*embryos (Fig. 5A and Movie S2). This result suggests that *pdgfab* is specifically required for the migration of sclerotome-derived cells to the medial trunk, but not to the peripheral fin fold. Consistent with these findings, *pdgfab^-/-^*embryos displayed substantially reduced staining of both *nkx3.1* and *nkx3.2* in sclerotome-derived cells around the notochord at 24 hpf (Fig. 5B). Similarly, the expression of the tenocyte marker *scxa* was also greatly reduced in *pdgfab^-/-^* fish at 48 hpf, whereas the expression of the fin mesenchymal cell marker *fbln1* remained largely unaffected (Fig. 5B). To confirm these observations, we quantified the number of tenocytes (Fig. 5C) and fin mesenchymal cells (Fig. 5D) at 56 and 48 hpf, respectively. Consistent with marker analysis, the number of tenocytes was significantly reduced in a dose-dependent manner in fish carrying different copies of the wild-type *pdgfab* allele, with averages decreasing from 19 per myotendinous junction in *pdgfab^+/+^* to 15 in *pdgfab^+/-^* and 10 in *pdgfab^-/-^* fish (Fig. 5C). By contrast, there was no significant difference in the number of fin mesenchymal cells among different genotypes (Fig. 5D). Notably, *pdgfab^+/-^* embryos displayed a mild migration phenotype (Fig. 5B-C), suggesting that the level of Pdgfab ligand is critical for the normal migration of sclerotome-derived cells. Together, our results suggest that the Pdgfab ligand primarily controls the medial migration of sclerotome-derived cells toward the trunk compartment, but not toward the fin fold. This region-specific role of *pdgfab* correlates well with its expression pattern shown above.

### Pdgfab acts as a chemoattractant in guiding the migration of sclerotome-derived cells

Next, we tested whether Pdgfab functions as a chemoattractant to guide the migration of Pdgfra-expressing cells from the sclerotome. To generate embryos with sparse *pdgfab*- expressing cells, we injected *pdgfab^-/-^*mutants with a low dose of the *hsp:pdgfab-2A-mCherry* plasmid (Fig. 6A). Injected embryos, along with uninjected controls, were heat shocked at 18 hpf and co-stained for *nkx3.1* and *mCherry* probes at 24 hpf (Fig. 6A). As shown in Fig. 5B, in *pdgfab^-/-^* controls, cells failed to migrate out of the sclerotome domains, resulting in no *nkx3.1^+^*cells around the notochord at 24 hpf (Fig. 6B). By contrast, injected *pdgfab^-/-^*embryos consistently exhibited strong migration of *nkx3.1^+^* cells toward *pdgfab*-expressing cells (*mCherry*^+^) (Fig. 6B). We focused our analysis on embryos containing isolated *mCherry*^+^ cells or cell clusters. Strikingly, *pdgfab* expression in individual muscle cells (93%, 27/29 cases), notochord cells (100%, 16/16 cases), or cells in the spinal cord (83%, 10/12 cases) effectively attracted the migration of *nkx3.1^+^*cells (Fig. 6B-C). By contrast, *pdgfab* expression in skin cells or cells in the dorsal spinal cord failed to trigger a robust migratory response of *nkx3.1^+^*cells from the ventral sclerotome domain (Fig. S2), suggesting that *pdgfab*^+^ cells need to be physically close to induce the migration of *pdgfra*-expressing cells. Interestingly, when *pdgfab* was ectopically expressed along the yolk extension, it induced abnormal ventral migration of *nkx3.1*^+^ cells (86%, 19/22 cases), resulting in a close juxtaposition of *nkx3.1^+^*cells and the *mCherry^+^* yolk epithelial cells (Fig. 6B-C). However, *pdgfab* expression in the ventral edge of the yolk extension failed to attract *nkx3.1^+^* cells ventrally (Fig. S2), likely due to the physical distance. To further characterize all cases with obvious chemoattraction (Fig. 6C), we quantified the extent of migration of *nkx3.1*^+^ cells from the ventral edge of the somite (Fig. 6D). Compared to somites in *pdgfab^-/-^* embryos, *pdgfab* expression in muscles, notochord, or the spinal cord significantly rescued the migration of *nkx3.1*^+^ cells toward the notochord to a similar extent (Fig. 6E). By contrast, *pdgfab* expression in the yolk extension resulted in the migration of *nkx3.1*^+^ cells in the opposite direction (Fig. 6E). Together, our results demonstrate that Pdgfab functions as a chemoattractant in vivo, guiding the migration of Pdgfra-expressing sclerotome-derived cells.

**Figure 6:**
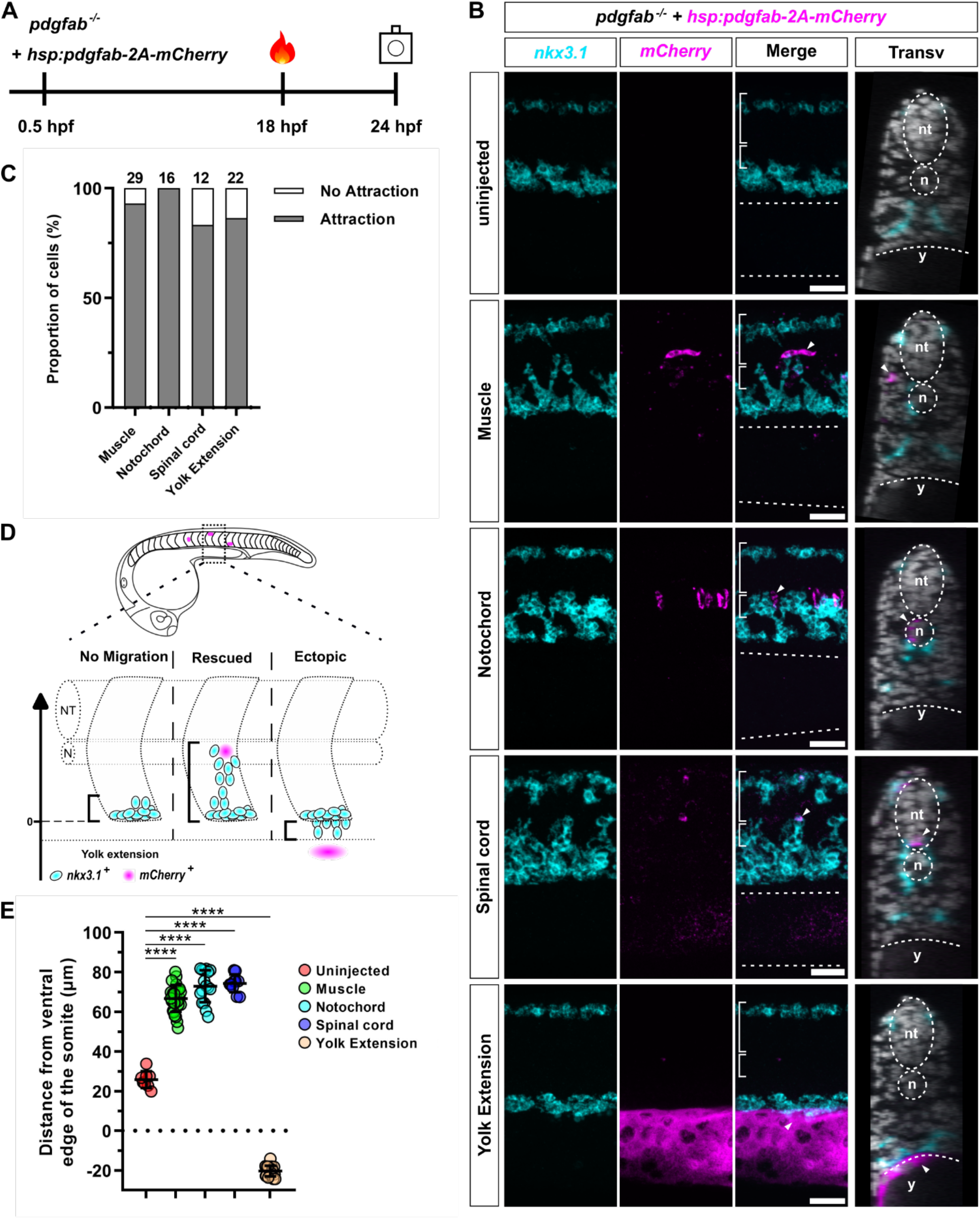
***pdgfab* acts as a chemoattractant for sclerotome-derived cells.** (A) Schematics of the experimental design. *pdgfab^-/-^*embryos were injected with the *hsp:pdgfab-2A-mCherry* plasmid at a low dose at the one-cell stage, heat shocked at 18 hpf, and double stained using *nkx3.1* and *mCherry* probes at 24 hpf. (B) Examples of *nkx3.1* (cyan) and *mCherry* (magenta) expression in uninjected and injected *pdgfab^-/-^* mutant embryos at 24 hpf. In sagittal views, the neural tube (long brackets), notochord (short brackets), and yolk extension (dotted lines) are labeled. In transverse views, the neural tube (nt), notochord (n), and yolk extension (y) are marked by dotted lines, and nuclear staining with Draq5 is shown. In uninjected controls (top), *nkx3.1*-expressing cells in the ventral sclerotome fail to migrate out to surround the notochord. By contrast, sparse expression of *hsp:pdgfab-2A-mCherry* in muscle, notochord, or spinal cord cells (*mCherry^+^*) substantially restores the migration of *nkx3.1*^+^ sclerotome-derived cells toward the notochord. Expression of *hsp:pdgfab-2A-mCherry* in the yolk extension leads to the ventral migration of *nkx3.1*^+^ cells toward the yolk. Localized *mCherry^+^ pdgfab*-expressing cells that attract *nkx3.1*^+^ sclerotome-derived cells are indicated by arrowheads. *n* = 10 (uninjected) and 80 (injected) embryos. (C) Quantification of the experiment shown in (B). The proportion of *mCherry^+^* cells (or cell clusters) that display attraction of *nkx3.1*^+^ cells is shown for the tissues depicted in (B). The number of cases for each tissue is indicated above each bar. (D) Schematics of the quantification of the migration extent of *nkx3.1*^+^ cells in the experiment showed in (B). We measured the distance between the ventral border of the somite (position = 0) and the leading edge of the *nkx3.1*^+^ cells. In uninjected *pdgfab^-/-^* mutants (“no migration”), the small distance represents the height of the ventral sclerotome domain. In *hsp:pdgfab-2A-mCherry*-injected *pdgfab^-/-^* embryos, *nkx3.1*^+^ cells derived from the ventral sclerotome can migrate toward the notochord (“rescued”, positive migration) or the yolk extension (“ectopic”, negative migration). (E) Quantification of the migration extent of *nkx3.1*^+^ cells as depicted in (D) for each of the scenarios shown in (B). Only cases where attraction was observed are plotted here, except in the uninjected animals. *n* = 8 (uninjected), 27 (muscle), 16 (notochord), 10 (spinal cord), and 19 (yolk extension) cases from 10 (uninjected) and 80 (injected) embryos. Statistics: One-way ANOVA with Dunnett’s multiple comparison test. Asterisks representation: *p* < 0.0001 (****). Scale bars: 50 μm.

## DISCUSSION

In this study, we combine in vivo imaging with gain- and loss-of-function approaches to investigate how fibroblast precursors are guided to specific locations in the developing zebrafish embryo. Our experiments reveal three key findings (Fig. 7). First, Pdgfra functions cell-autonomously in the migration of sclerotome-derived fibroblast precursors. Second, *pdgfab* is specifically required for the medial migration of sclerotome-derived cells. Third, localized Pdgfab expression acts as a chemoattractant in directing the migration of sclerotome-derived cells. Together, these results demonstrate that Pdgfra signaling mediates the chemoattraction of sclerotome-derived fibroblast precursors toward Pdgfab-expressing regions.

**Figure 7:**
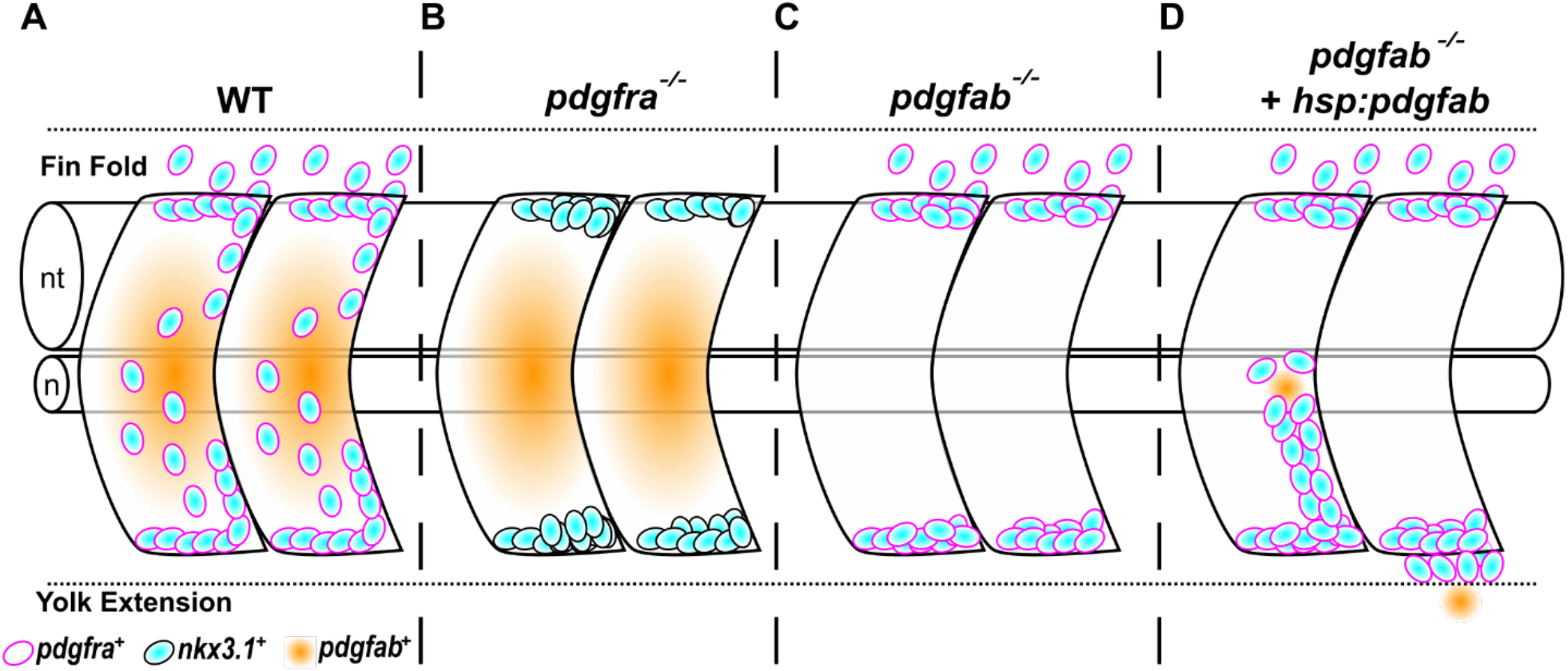
The migration of sclerotome-derived cells is regulated by chemoattraction mediated by Pdgfab/Pdgfra signaling. (A) In wild-type embryos, *nkx3.1^+^* (cyan) sclerotome-derived cells express the receptor Pdgfra (magenta), while the medial somites express the ligand Pdgfab (orange) in response to Hh signaling. *pdgfra^+^* sclerotome-derived cells undergo extensive migration toward both the notochord in the medial trunk and the fin fold. (B) In *pdgfra^-/-^* mutants, *nkx3.1^+^* sclerotome-derived cells fail to migrate in any direction and remain in their original location in the ventral and dorsal parts of the somites. (C) In *pdgfab^-/-^* embryos, *pdgfra^+^* sclerotome-derived cells fail to migrate toward the notochord, although migration toward the fin fold remains unaffected. (D) In *pdgfab^-/-^*embryos with mosaic *pdgfab* expression, *nkx3.1^+^* sclerotome cells can migrate toward the ectopic *pdgfab-*expressing cells, either medially around the notochord or ventrally toward the yolk extension. When the *pdgfab* source is located near or in the notochord, it can completely rescue the migration of *nkx3.1^+^* sclerotome cells toward the notochord. These diagrams depict a 2-somite region located above the yolk extension of a zebrafish embryo at 48 hpf. nt: neural tube; n: notochord.

### Pdgfra is required cell-autonomously for the migration of sclerotome-derived fibroblast precursors

Previous studies in mice and chicks have shown that loss of PDGFRA function leads to various axial skeleton defects, suggesting compromised sclerotome development (Pickett et al., 2008; Soriano, 1997). However, how PDGFRA signaling regulates the development of sclerotome-derived cells in vivo remains unclear. In our study, we provide three lines of evidence demonstrating that *pdgfra* is essential for the migration of sclerotome-derived cells in zebrafish (Fig. 7A-B). First, *pdgfra* is specifically enriched in the sclerotome and its derivatives. Second, loss-of-function experiments, using either the *pdgfra* gene-trap mutant or overexpression of dominant-negative PdgfraΔTK-EGFP, result in a complete disruption of sclerotome-derived cell migration, accompanied by the loss of sclerotome derivatives such as tenocytes and fin mesenchymal cells. Third, mosaic inhibition of Pdgfra activity by membrane-tethered PdgfraΔK-EGFP specifically blocks the migration of PdgfraΔK-EGFP-expressing sclerotome cells but not non-expressing cells, indicating that Pdgfra functions in a cell-autonomous manner.

The *pdgfra^mRFP^* gene-trap allele is predicted to produce a secreted, truncated Pdgfra receptor fused to mRFP (PdgfraΔTK-mRFP), lacking both the transmembrane and kinase domains. With an intact ligand-binding extracellular domain but no signaling capability, PdgfraΔTK-mRFP likely functions as a decoy receptor, sequestering Pdgf ligands in the extracellular space. Similar decoy receptors, such as interleukin-1 type II receptor (IL-1RII) and soluble vascular endothelial growth factor receptor-1 (sVEGFR-1), have been shown to inhibit their respective signaling pathways (Colotta et al., 1993; Ebos et al., 2004). Although naturally occurring secreted short forms of PDGFRA have not been reported, muscle-resident fibro/adipogenic progenitors express a short isoform produced by an internal polyadenylation site (Mueller et al., 2016). This isoform functions as a membrane-tethered decoy receptor, attenuating PDGF signaling during muscle regeneration (Mueller et al., 2016). Consistent with the dominant nature of a decoy receptor, heterozygous *pdgfra^mRFP/+^* embryos exhibit reduced migration and differentiation of sclerotome-derived cells compared to their wild-type siblings. Interestingly, the mRFP signal from PdgfraΔTK-mRFP is undetectable in the extracellular space and only gradually appears later in the endothelial scavenger cells (Campbell et al., 2018). Similar vascular localization has been observed in other gene-trap lines targeting genes with secreted signals at their 5’ end (Clark et al., 2011), suggesting that these secreted truncated proteins are captured by endothelial scavenger cells.

Consistent with our findings, *pdgfra* has been recently shown to be required for the development of stromal reticular cells in zebrafish (Murayama et al., 2023). These cells, which support hematopoietic stem cells, are derived from the ventral sclerotome domain in the tail (Ma et al., 2023; Murayama et al., 2015). Using morpholino knockdown experiments, Murayama et al. show that *pdgfra* is required for the ventral migration of sclerotome cells to give rise to stromal reticular cells and fin mesenchymal cells, but is not necessary for dorsal migration toward the notochord (Murayama et al., 2023). This result contrasts with our findings that *pdgfra* is critical for the migration of sclerotome-derived cells in all directions, including toward both the notochord and the fin fold. This discrepancy may be due to the stronger phenotype caused by the dominant-negative nature of the *pdgfra* gene-trap mutant compared to the morpholino knockdown. Additionally, it is possible that the ventral migration of sclerotome cells toward the fin fold requires a higher level of Pdgfra activation, making it more sensitive to pathway inhibition.

### Hh signaling-dependent *pdgfab* is required for sclerotome-derived cell migration toward the medial trunk

Previous studies in chick suggest that Hh signaling is essential for sclerotome development (Cairns et al., 2008). Our group has shown that in zebrafish, Hh signaling is dispensable for the induction of sclerotome domains but is essential for the migration and maintenance of marker expression in sclerotome-derived cells around the notochord (Ma et al., 2018). Here, we show that *pdgfab* exhibits partially overlapping expression in the medial somite with other Hh target genes such as *ptc2* and *myod1*. Inhibition of Hh signaling by cyclopamine results in a complete loss of *pdgfab* expression in the somites, accompanied by impaired migration of sclerotome-derived cells toward the notochord. This migration phenotype is recapitulated in *pdgfab* mutants, suggesting that Hh signaling indirectly regulates the migration of sclerotome-derived cells by controlling *pdgfab* expression.

However, we cannot rule out the possibility of a parallel mechanism, in which Hh signaling- dependent adaxial cell specification and lateral migration (Daggett et al., 2007; Ono et al., 2015) serve as prerequisites for triggering the migration of sclerotome cells toward the notochord (Stickney et al., 2000). A comparison of migration phenotypes between the *pdgfra* gene-trap line and *pdgfab* mutants reveals two key differences. First, *pdgfra* mutants show defects of sclerotome cell migration in all directions, leading to the loss of both tenocytes and fin mesenchymal cells (Fig. 7B). By contrast, *pdgfab* mutants specifically show impaired sclerotome cell migration toward the notochord, while the migration of fin mesenchymal cell precursors into the fin fold remains largely unaffected (Fig. 7C). We predict that sclerotome-derived cell migration is orchestrated by multiple Pdgfra ligands expressed in distinct regions. Based on our expression analysis, *pdgfc* is the only ligand expressed in the distal edge of the fin fold, the target area for fin mesenchymal cell precursors. Thus, it is plausible that Pdgfab acts as a chemoattractant to mediate migration toward the notochord region, while Pdgfc plays a similar role in the fin fold. Interestingly, we did not observe enhanced migration toward the fin fold in the absence of *pdgfab*, suggesting that additional mechanisms, such as contact inhibition reported in neural crest cells (Bahm et al., 2017), may regulate the number of fibroblast precursors in target regions.

Another difference between *pdgfra* and *pdgfab* mutants is that *pdgfra* embryos exhibit a more severe phenotype in the migration of sclerotome-derived cells toward the notochord, resulting in a greater reduction in tenocyte number. The expression of both *pdgfaa* and *pdgfc* in the medial trunk region suggests that these ligands may function redundantly with Pdgfab to mediate the migration of sclerotome-derived cells in this area. Similar functional redundancy has been observed between *pdgfaa* and *pdgfab* in zebrafish during pharyngeal arch artery morphogenesis (Mao et al., 2019) and primitive erythropoiesis (Mao et al., 2023).

### Sclerotome-derived cell migration is regulated by chemoattraction through Pdgfab/Pdgfra signaling

Chemoattraction by PDGF ligands has been first described in vitro in experiments with smooth muscle cells and fibroblasts (Grotendorst et al., 1981; Seppa et al., 1982). In vivo studies in *Drosophila* have further demonstrated a chemoattractive role of the homologous PDGF signaling pathway in border cell migration (Duchek et al., 2001; McDonald et al., 2003). During early oogenesis in *Drosophila*, the oocyte expresses PDGF/VEGF-related factor 1 (PVF1), which acts as a chemoattractant to guide the migration of border cells expressing PVR, a receptor tyrosine kinase related to the mammalian PDGF and VEGF receptors (Duchek et al., 2001; McDonald et al., 2003). Similar chemoattraction models have been proposed to explain the directed migration of both cardiomyocytes and neural crest cells in vertebrates, where PDGFRA-expressing cells migrate toward PDGFA ligand-expressing target areas (Bahm et al., 2017; Bloomekatz et al., 2017; Eberhart et al., 2008). Indeed, PDGFA-coated beads have been shown to direct the migration of PDGFRA-expressing neural crest cells in vivo in both zebrafish and mice (Eberhart et al., 2008; Kawakami et al., 2011), as well as in vitro in xenopus explants (Bahm et al., 2017). These findings suggest that PDGFA/PDGFRA signaling-mediated chemoattraction may be a conserved mechanism in regulating directed cell migration in various developmental contexts.

Previous studies have suggested a role for PDGFRA signaling in regulating the migration of sclerotome-derived cells in mice (Pickett et al., 2008; Soriano, 1997; Tallquist et al., 2000); however, this function has been inferred rather than directly demonstrated due to technical limitations. In our current study, we provide in vivo evidence that Pdgfab acts as an instructive cue, likely functioning as a chemoattractant to guide the migration of Pdgfra- expressing fibroblast precursors in zebrafish. First, ubiquitous overexpression of Pdgfab completely abolishes the migration of sclerotome-derived cells. This suggests an instructive rather than permissive role for Pdgfab, as uniform distribution of the signal eliminates the directional cues necessary for guided migration. Second, in the absence of endogenous Pdgfab, mosaic expression of Pdgfab in medial cells, such as muscles, notochord, and ventral spinal cord cells, is sufficient to rescue the localized migration of sclerotome-derived cells to surround the notochord (Fig. 7D). Interestingly, although the notochord does not normally express *pdgfab*, it efficiently attracts sclerotome-derived cells, likely due to its position as their final migration destination (Ma et al., 2018). This result supports a model in which the notochord can secrete functional Pdgfab protein, suggesting that the specific cell type producing the ligand is less critical than where sclerotome-derived cells encountered the protein. Our mosaic expression experiments further reveal that the secreted Pdgfab protein has a limited diffusion range, as *pdgfab* expression in distant tissues, such as the skin, dorsal spinal cord, or dorsal muscle fibers, is insufficient to trigger the migration of cells from the ventral sclerotome domain. This localized effect underscores the importance of spatially restricted Pdgfab expression for proper cell guidance. Strikingly, ectopic Pdgfab expression in the yolk extension is sufficient to redirect sclerotome-derived cells to undergo abnormal ventral migration (Fig. 7D). This result mirrors previous findings where ectopic PVF1 expression similarly redirects border cell migration in *Drosophila* (McDonald et al., 2003).

Collectively, our findings strongly indicate that Pdgfa/Pdgfra signaling-mediated chemoattraction guides the migration of sclerotome-derived cells in zebrafish. Given that *Pdgfa* is expressed in the myotome in mice (Orr-Urtreger & Lonai, 1992; Tallquist et al., 2000) and *Pdgfra* is expressed in the sclerotome in both mice and humans (Loh et al., 2016; Orr-Urtreger & Lonai, 1992), we predict that the migration of sclerotome-derived cells in mammals is likely guided by a similar chemotactic mechanism.

In summary, our zebrafish studies demonstrate that Pdgfab functions as a localized chemoattractant cue to guide the migration of Pdgfra-expressing fibroblast precursors from the sclerotome (Fig. 7). This work provides a framework for studying the development and diversification of distinct fibroblast subtypes during development. Our findings also suggest that similar PDGFA/PDGFRA signaling mechanisms may be employed in other contexts, such as recruiting fibroblasts during wound healing or regulating cell migration in pathological conditions like cancer metastasis (Guérit et al., 2021). Targeting PDGFA/PDGFRA signaling could represent a therapeutic opportunity to treat fibrosis-related disorders and cancer.

## MATERIALS AND METHODS

### Ethics statement

All animal experiments were conducted in accordance with the principles outlined in the current guidelines of the Canadian Council on Animal Care. All protocols were approved by the Animal Care Committee at the University of Calgary (#AC21-0102).

### Zebrafish strains

Zebrafish strains used were raised under standard zebrafish welfare conditions (Bradford et al., 2022). The following lines were utilized: *pdgfra^smh1300Gt^*(referred to as *pdgfra^mRFP^*) (El-Rass et al., 2017), *pdgfab^sa11805^* (Kettleborough et al., 2013), *Tg(hsp:pdgfab- 2A-mCherry,cryaa:CFP)ca118* (referred to as *hsp:pdgfab-2A-mCherry*), *Tg(kdrl:EGFP)la163* (Choi et al., 2007), *TgBAC(nkx3.1:Gal4)ca101* (Ma et al., 2018), and *Tg(UAS:NTR- mCherry)c264* (Davison et al., 2007). Fluorescent transgenic lines were screened by the expression of the respective fluorescent marker. The *pdgfra^mRFP^*line was maintained as heterozygotes, as homozygous mutants are not viable beyond 14 dpf (El-Rass et al., 2017). The *pdgfab^sa11805^* line was maintained as both heterozygotes and homozygotes.

### Mutant genotyping

The genotype of embryos carrying the *pdgfra^mRFP^* or *pdgfab^sa11805^* allele was determined using conventional PCR with allele-specific primers. Genomic DNA was extracted using the HOTSHOT protocol. Briefly, tissue from embryos or fin clips was dissociated in 50 mM NaOH at 95°C for 20 minutes, following by cooling on ice and neutralization with 1 M Tris- HCl (pH = 7.5). To identify the *pdgfra^mRFP^* allele, the primers pdgfra-g-F3 (5’- TCGGACCACGAGTCTACATA-3’) and mRFP-1 (5’-GAGCCCTCCATGCGCACCTTGAA-3’) were used with an annealing temperature of 61°C, producing a single 805 bp band in homozygous mutant embryos. For wild-type embryos, the primers pdgfra-g-F3 and pdgfra-g2- R (5’-TGTGTCAACGCCACAACCTA-3’) were used with an annealing temperature of 61°C, yielding a single 575 bp band. PCRs from heterozygous embryos produce both corresponding bands.

The *pdgfab^sa11805^* allele has a 2 bp change from 5’-AT<u>C</u>TA<u>C</u>-3’ to 5’-ATtTA<u>a</u>-3’ at positions 333 and 336 of the coding sequence, resulting in a substitution of tyrosine (Y) at amino acid 112 in the wild-type allele with a premature stop codon in *pdgfab^sa11805^*. We designed a one-step PCR assay (J. Chen & Schedl, 2021) to distinguish the *pdgfab^sa11805^*allele from the wild-type allele. To identify the *pdgfab^sa11805^*allele, the primers Pdgfab_Geno3_Fmut (5’-CAAGACCCGGACGGTGCTT-3’) and Pdgfab_Geno2_R (5’- CAAACACACAAACCCGTGAC-3’) were used with an annealing temperature of 62°C, producing a single 171 bp band in homozygous mutant embryos. For wild-type embryos, the primers Pdgfab_Geno4_Fwt (5’-CAAGACCCGGACGGTGCTC-3’) and Pdgfab_Geno2_R were used with an annealing temperature of 62°C, also resulting in a single 171 bp band.

PCRs from heterozygous embryos produce both corresponding bands.

### Generation of expression constructs

All expression constructs were assembled following the SLiCE protocol (Zhang et al., 2012). The *hsp70:pdgfab-2A-mCherry* plasmid was generated by replacing the *pdgfaa* sequence in the *hsp70:pdgfaa-2A-mCherry* plasmid (Bloomekatz et al., 2017) with the *pdgfab* coding sequence.

To generate the *UAS:pdgfraΔK-EGFP* plasmid, *UAS:pdgfra* was first constructed by replacing *NTR-mCherry* in the *UAS:NTR-mCherry* plasmid with the full-length *pdgfra* coding sequence (ENSDART00000103510.5). The cytosolic domain of *pdgfra* at position 2838-4397 of the coding sequence was then replaced with a linker sequence (5’- AGTCTCGGACCTGGACTCGGATCCGGA-3’) and an ATG-less *EGFP* sequence.

The *hsp:pdgfraΔTK-EGFP* plasmid was generated in two steps: First, *pdgfraΔK-EGFP* was inserted into *hsp70:EGFP* to replace *EGFP*. Second, the transmembrane domain coding sequence was removed using a PCR strategy (Hansson et al., 2008; Krishnamurthy et al., 2015).

### Generation of transgenic lines

The *Tg(hsp:pdgfab-2A-mCherry,cryaa:CFP)* line was generated using the *hsp70:pdgfab-2A-mCherry* plasmid, which carries *cryaa:CFP* as a transgenesis marker. To generate a stable line, 40 pg of the *hsp70:pdgfab-2A-mCherry* plasmid was co-injected with 40 pg of *tol2* mRNA into one-cell stage wild-type embryos. The injected embryos were screened at 3 dpf for *cryaa:CFP* expression in the lens. Positive embryos were raised to adulthood as potential founders. Stable transgenic lines were established by screening for *cryaa:CFP* expression in F1 embryos from these injected founders.

### Embryo injections

For most plasmid injections, 40 pg of plasmid DNA was co-injected with 40 pg of *tol2* transposase mRNA into embryos at the one-cell stage. For mosaic experiments with *UAS:pdgfraτιK-EGFP*, 10 pg of the plasmid was injected. For mosaic experiments with *hsp:pdgfab-2A-mCherry*, 2 pg of the plasmid was injected without *tol2* transposase mRNA. For *Cre* expression, 20 pg of *Cre* mRNA was injected into one-cell stage embryos obtained from intercrosses of *pdgfra^mRFP/+^* fish. To label scavenger endothelial cells, 1 nL of a 2 mg/mL solution of Alexa Fluor 647-conjugated ovalbumin (Invitrogen, O34784) was injected into the extracellular space of embryos at 4 hpf.

### Heat shock induction

To induce heat shock promoter-driven genes, the embryos were transferred into a 2 mL tube and placed in a heat block set at 37°C for 30 minutes, at the desired developmental stage. Afterward, the embryos were transferred to E3 fish water and allowed to continue developing at 28.5°C until fixation.

### Time-lapse imaging

Time-lapse imaging of zebrafish embryos was conducted using an Olympus FV1200 confocal microscope. Embryos at the desired developmental stage were anesthetized with tricaine and mounted in a 35 mm glass-bottom imaging dish. A small volume of E3 fish water containing phenylthiourea (PTU) and tricaine was carefully added around the agarose to prevent pigment development and minimize embryo movement. The trunk region above the yolk extension was imaged, with Z-stack images captured at intervals specified in each figure. The images were then compiled and processed using Fiji software (Schindelin et al., 2012).

### In situ hybridization

Whole mount in situ hybridization were performed following standard protocols (Thisse & Thisse, 2008). The RNA probes used were: *fbln1*, *fras1*, *GFP*, *mCherry*, *mRFP*, *myod1*, *nkx3.1*, *nkx3.2*, *pax1a*, *pax9*, *pdgfaa*, *pdgfab*, *pdgfc*, *pdgfra*, *pdgfra5’*, *pdgfra3’*, *prelp*, *ptc2*, and *scxa.* Double fluorescent in situ hybridizations were perform combining digoxigenin (DIG) and dinitrophenyl (DNP) labeled probes. Draq5 (1:10,000, Biostatus, DR50050) was used for nuclear staining. To obtain transverse views, stained embryos were manually sectioned using vibratome steel blades.

### Drug treatment

Embryos at 5 hpf were treated in 100 μM cyclopamine (Cedarlane Labs, C988400-50) in E3 fish water. Control embryos were treated with 1% DMSO in E3 fish water. The treated embryos were maintained at 28°C until 24 hpf for analysis.

### Quantifications

To quantify *nkx3.1*-stained area, stained embryos were imaged, and a region of interest comprising 8 somites above the yolk extension was defined. The image was then converted to black and white, a threshold was set, and the coverage ratio was calculated. These values were subsequently plotted on graphs.

For tenocyte counting, the *nkx3.1:Gal4; UAS:NTR-mCherry* reporter was used in the appropriate genetic background at the desired developmental stage. The number of tenocytes in three myotendinous junctions per animal was counted, and the average number per MTJ for each fish was then plotted on graphs.

To quantify fin mesenchymal cells, anesthetised animals at the desired developmental stage were observed under an inverted Zeiss microscope with a 20x objective. Fin mesenchymal cells, identified by their characteristic morphology, were counted both dorsally and ventrally in an area spanning 8 somites posterior to the end of the yolk extension. The counts were summed for each individual fish and plotted on graphs.

For the chemoattraction experiments, the extent of migration of the *nkx3.1^+^* cells was determined by measuring the distance between the ventral edge of the somite and the leading *nkx3.1^+^* cell beneath an *mCherry^+^* cell. The measurements were performed using Fiji software (Schindelin et al., 2012).

### Statistical analysis

All the graphs were generated using the GraphPad Prism software. Data were plotted as mean ± SD. Significance was calculated using these statistic tests: One-way ANOVA with Tukey’s multiple comparison test (Fig. 1E, 2C-D, and 5C-D), Unpaired t test with Welch’s correction (Fig. 3E), and One-way ANOVA with Dunnett’s multiple comparison test (Fig. 6E). *p* values: *p* < 0.0001 (****), *p* < 0.001 (***), *p* < 0.01 (**), *p* < 0.05 (*), and *p* > 0.05 (ns, not significant).

## ACKNOWLEDGMENTS

We thank the zebrafish community for sharing fish lines and reagents, particularly Joshua Bloomekatz, Alexa Burger, Holger Knaut, Xiao-Yan Wen, and Zebrafish International Resource Center (ZIRC). We are also grateful to Sarah Childs for sharing transgenic lines and providing critical input on this project, as well as to the members of the Childs and Huang laboratories for their valuable discussions.

## Author Contributions

APG and MCC contributed to conception and design of the study. APG, NBC, FT, CLG and MD carried out experimental work. APG and MCC wrote the first draft of the manuscript. All authors contributed to manuscript revision, read, and approved the submitted version. Funding and Resources: MCC; Supervision: APG and MCC.

---

**Figure S1:**
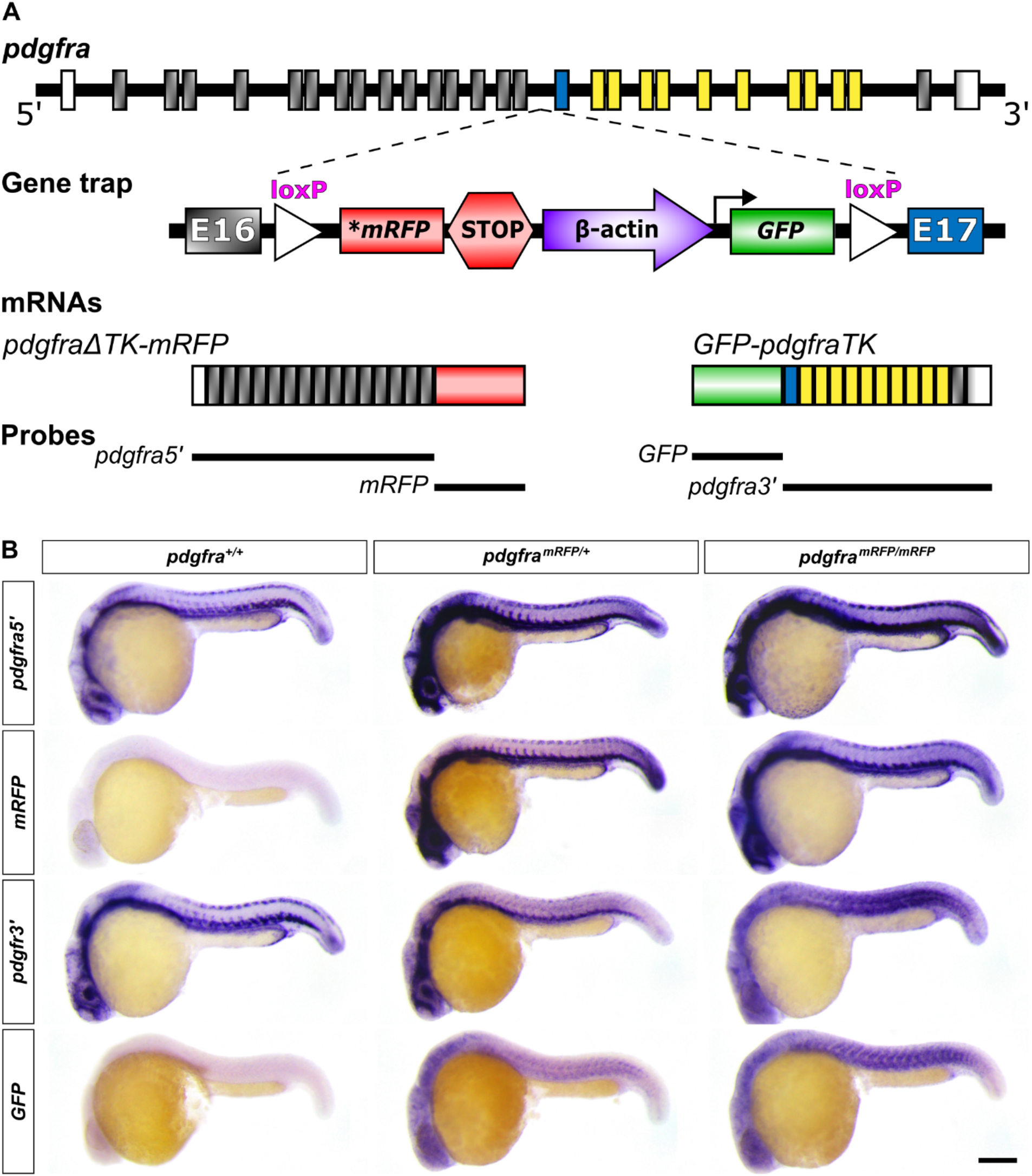
Characterization of the *pdgfra* gene-trap line. (A) Schematics of the *pdgfra* gene-trap mutant line. The exons of the *pdgfra* gene are color-coded: gray for extracellular coding exons, blue for the transmembrane coding exon, yellow for intracellular coding exons, and white for non-coding exons. This *pdgfra* gene-trap line is predicted to produce two mRNAs: the extracellular domain of *pdgfra* fused with *mRFP* (*pdgfraΔTK-mRFP*) under the control of endogenous *pdgfra* promoter, and GFP fused with the transmembrane and cytoplasmic kinase domains of *pdgfra* (*GFP-pdgfraTK*) under the control of the *β-actin* promoter. The locations of different region-specific probes used to characterize this mutant are shown as black lines aligned underneath the corresponding mRNA. (B) Expression analysis of the mRNAs produced from the *pdgfra* gene-trap line. *pdgfra^+/+^*, *pdgfra^mRFP/+^*, and *pdgfra^mRFP/mRFP^* embryos were stained at 24 hpf using two probes against the endogenous *pdgfra* sequences (*pdgfra5’* and *pdgfra3’*) and two probes against the exogenous inserts (*mRFP* and *GFP*). Only *pdgfra^mRFP/+^* and *pdgfra^mRFP/mRFP^* embryos express *mRFP* and *GFP*. Scale bar: 100 μm.

**Figure S2:**
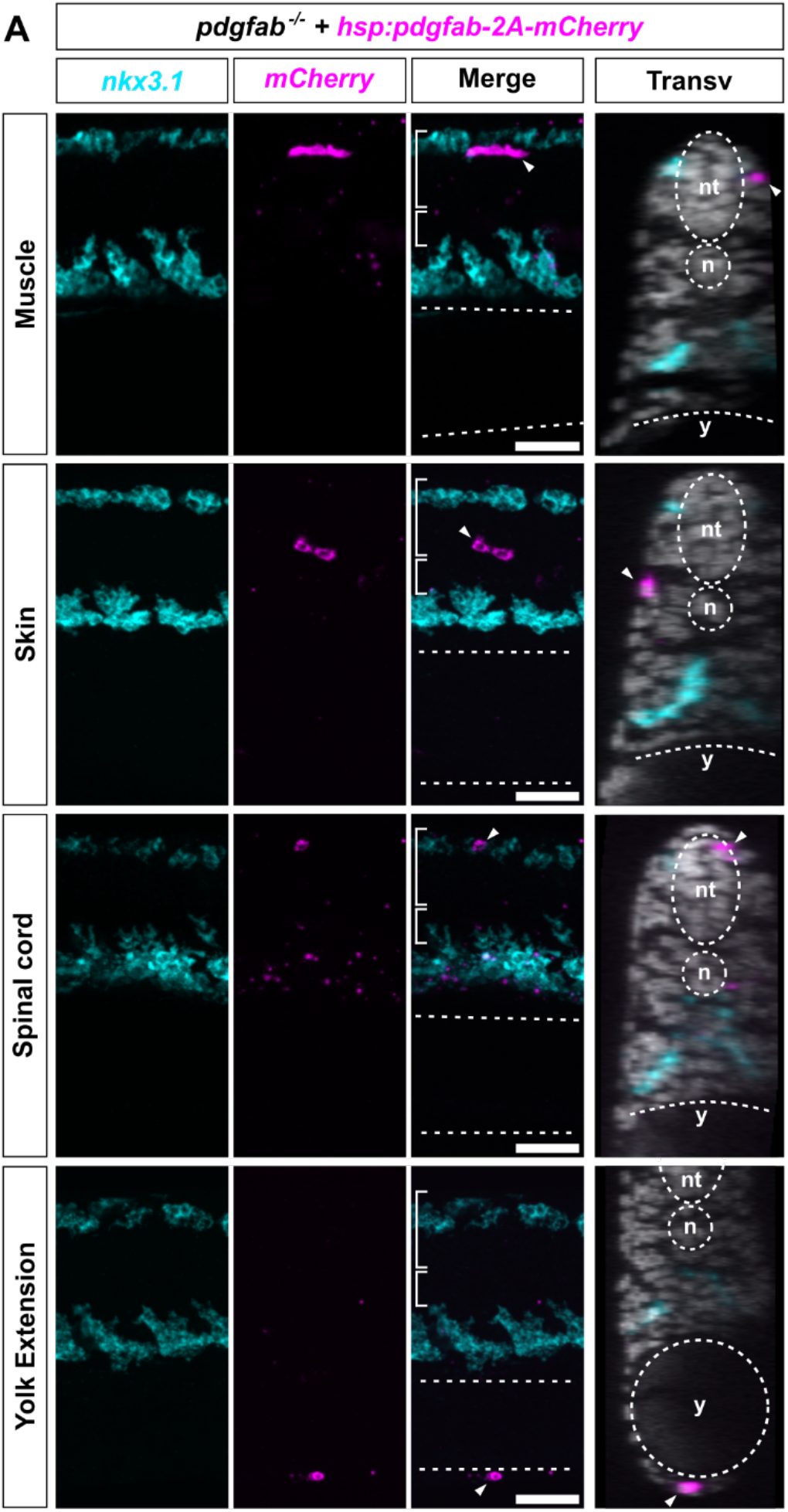
Examples of cases showing no chemoattraction in response to mosaic *pdgfab* expression. *pdgfab^-/-^*embryos were injected with the *hsp:pdgfab-2A-mCherry* plasmid at a low dose at the one-cell stage, heat shocked at 18 hpf, and double stained using *nkx3.1* (cyan) and *mCherry* (magenta) probes at 24 hpf. Examples of cases showing no chemoattraction are shown here. In sagittal views, the neural tube (long brackets), notochord (short brackets), and yolk extension (dotted lines) are labeled. In transverse views, the neural tube (nt), notochord (n), and yolk extension (y) are marked by dotted lines, and nuclear staining with Draq5 is shown. *pdgfab* expression in dorsal muscle fibers, skin cells, dorsal spinal cord cells, or yolk cells at the ventral edge of the yolk extension does not induce migration of *nkx3.1^+^* cells from the ventral sclerotome domain. *mCherry^+^ pdgfab*-expressing cells are indicated by arrowheads. *n* = 2 (muscle), 2 (spinal cord), and 3 (yolk extension) cases from 80 (injected) embryos. Scale bars: 50 μm.

**Movie S1: Time-lapse imaging of sclerotome-derived cells in wild-type and *pdgfra* gene-trap mutant backgrounds.** Embryos at 22 hpf were imaged every 17 minutes for 14 hours. In wild-type siblings, sclerotome-derived cells above the yolk extension migrate from the ventral and dorsal sclerotome domains toward the notochord. In contrast, heterozygous and homozygous individuals exhibit reduced migration in a dose-dependent manner. Snapshots from this time-lapse are shown in Figure 2A. Time stamps are indicated in the hh:mm format. Scale bar: 100 μm.

**Movie S2: Time-lapse imaging of sclerotome-derived cells in wild-type and *pdgfab* mutant backgrounds.** Embryos at 24 hpf were imaged every 20 minutes for 10 hours. In wild-type siblings, sclerotome-derived cells migrate from the ventral and dorsal sclerotome domains toward the notochord. *pdgfab^+/-^* individuals show slight migration defects, while *pdgfab^-/-^* mutants exhibit a more pronounced reduction in migration toward the notochord, though migration toward the fin folds remains largely unaffected. Snapshots from this time- lapse are shown in Figure 5A. Time stamps are indicated in the hh:mm format. Scale bar: 100 μm.

## REFERENCES

1. Andrae, J., Gallini, R., & Betsholtz, C. (2008). Role of platelet-derived growth factors in physiology and medicine. Genes & Development, 22(10), 1276–1312. 10.1101/gad.1653708

2. Bahm, I., Barriga, E. H., Frolov, A., Theveneau, E., Frankel, P., & Mayor, R. (2017). PDGF controls contact inhibition of locomotion by regulating N-cadherin during neural crest migration. Development, 144, 2456–2468. 10.1242/dev.147926

3. Bloomekatz, J., Singh, R., Prall, O. W., Dunn, A. C., Vaughan, M., Loo, C.-S., Harvey, R. P., & Yelon, D. (2017). Platelet-derived growth factor (PDGF) signaling directs cardiomyocyte movement toward the midline during heart tube assembly. eLife, 6, e21172. 10.7554/eLife.21172

4. Boström, H., Willetts, K., Pekny, M., Levéen, P., Lindahl, P., Hedstrand, H., Pekna, M., Hellström, M., Gebre-Medhin, S., Schalling, M., Nilsson, M., Kurland, S., Törnell, J., Heath, J. K., & Betsholtz, C. (1996). PDGF-A Signaling Is a Critical Event in Lung Alveolar Myofibroblast Development and Alveogenesis. Cell, 85(6), 863–873. 10.1016/S0092-8674(00)81270-2

5. Bradford, Y. M., Van Slyke, C. E., Ruzicka, L., Singer, A., Eagle, A., Fashena, D., Howe, D. G., Frazer, K., Martin, R., Paddock, H., Pich, C., Ramachandran, S., & Westerfield, M. (2022). Zebrafish information network, the knowledgebase for Danio rerio research. Genetics, 220(4), iyac016. 10.1093/genetics/iyac016

6. Cairns, D. M., Sato, M. E., Lee, P. G., Lassar, A. B., & Zeng, L. (2008). A gradient of Shh establishes mutually repressing somitic cell fates induced by Nkx3.2 and Pax3. Developmental Biology, 323(2), 152–165. 10.1016/j.ydbio.2008.08.024

7. Campbell, F., Bos, F. L., Sieber, S., Arias-Alpizar, G., Koch, B. E., Huwyler, J., Kros, A., & Bussmann, J. (2018). Directing Nanoparticle Biodistribution through Evasion and Exploitation of Stab2-Dependent Nanoparticle Uptake. ACS Nano, 12(3), 2138–2150. 10.1021/acsnano.7b06995

8. Chen, J., & Schedl, T. (2021). A simple one-step PCR assay for SNP detection. microPublication Biology, Jun 11. 10.17912/micropub.biology.000399

9. Chen, P.-H., Chen, X., & He, X. (2013). Platelet-derived growth factors and their receptors: Structural and functional perspectives. Biochimica et Biophysica Acta (BBA) - Proteins and Proteomics, 1834(10), 2176–2186. 10.1016/j.bbapap.2012.10.015

10. Choi, J., Dong, L., Ahn, J., Dao, D., Hammerschmidt, M., & Chen, J.-N. (2007). FoxH1 negatively modulates flk1 gene expression and vascular formation in zebrafish. Developmental Biology, 304(2), 735–744. 10.1016/j.ydbio.2007.01.023

11. Clark, K. J., Balciunas, D., Pogoda, H.-M., Ding, Y., Westcot, S. E., Bedell, V. M., Greenwood, T. M., Urban, M. D., Skuster, K. J., Petzold, A. M., Ni, J., Nielsen, A. L., Patowary, A., Scaria, V., Sivasubbu, S., Xu, X., Hammerschmidt, M., & Ekker, S. C. (2011). In vivo protein trapping produces a functional expression codex of the vertebrate proteome. Nature Methods, 8(6), 506–512. 10.1038/nmeth.1606

12. Collins, F. L., Roelofs, A. J., Symons, R. A., Kania, K., Campbell, E., Collie-duguid, E. S. R., Riemen, A. H. K., Clark, S. M., & De Bari, C. (2023). Taxonomy of fibroblasts and progenitors in the synovial joint at single-cell resolution. Annals of the Rheumatic Diseases, 82(3), 428–437. 10.1136/ard-2021-221682

13. Colotta, F., Re, F., Muzio, M., Bertini, R., Polentarutti, N., Sironi, M., Giri, J. G., Dower, S. K., Sims, J. E., & Mantovani, A. (1993). Interleukin-1 Type 11 Receptor: A Decoy Target for IL-1 That Is Regulated by IL-4. Science, 261, 472–475.

14. Concordet, J.-P., Lewis, K. E., Moore, J. W., Goodrich, L. V., Johnson, R. L., Scott, M. P., & Ingham, P. W. (1996). Spatial regulation of a zebrafish patched homologue reflects the roles of sonic hedgehog and protein kinase A in neural tube and somite patterning. Development, 122(9), 2835–2846. 10.1242/dev.122.9.2835

15. Daggett, D. F., Domingo, C. R., Currie, P. D., & Amacher, S. L. (2007). Control of morphogenetic cell movements in the early zebrafish myotome. Developmental Biology, 309(2), 169–179. 10.1016/j.ydbio.2007.06.008

16. Damm, E. W., & Winklbauer, R. (2011). PDGF-A controls mesoderm cell orientation and radial intercalation during Xenopus gastrulation. Development, 138(3), 565–575. 10.1242/dev.056903

17. Davison, J. M., Akitake, C. M., Goll, M. G., Rhee, J. M., Gosse, N., Baier, H., Halpern, M. E., Leach, S. D., & Parsons, M. J. (2007). Transactivation from Gal4-VP16 transgenic insertions for tissue-specific cell labeling and ablation in zebrafish. Developmental Biology, 304(2), 811–824. 10.1016/j.ydbio.2007.01.033

18. Draga, M., & Scaal, M. (2024). Building a vertebra: Development of the amniote sclerotome. Journal of Morphology, 285(1), e21665. 10.1002/jmor.21665

19. Driskell, R. R., & Watt, F. M. (2015). Understanding fibroblast heterogeneity in the skin. Trends in Cell Biology, 25(2), 92–99. 10.1016/j.tcb.2014.10.001

20. Duchek, P., Somogyi, K., Jékely, G., Beccari, S., & Rørth, P. (2001). Guidance of Cell Migration by the Drosophila PDGF/VEGF Receptor. Cell, 107(1), 17–26. 10.1016/S0092-8674(01)00502-5

21. Eberhart, J. K., He, X., Swartz, M. E., Yan, Y.-L., Song, H., Boling, T. C., Kunerth, A. K., Walker, M. B., Kimmel, C. B., & Postlethwait, J. H. (2008). MicroRNA Mirn140 modulates Pdgf signaling during palatogenesis. Nature Genetics, 40(3), 290–298. 10.1038/ng.82

22. Ebos, J. M. L., Bocci, G., Man, S., Thorpe, P. E., Hicklin, D. J., Zhou, D., Jia, X., & Kerbel, R. S. (2004). A Naturally Occurring Soluble Form of Vascular Endothelial Growth Factor Receptor 2 Detected in Mouse and Human Plasma. Molecular Cancer Research, 2(6), 315–326. 10.1158/1541-7786.315.2.6

23. El-Rass, S., Eisa-Beygi, S., Khong, E., Brand-Arzamendi, K., Mauro, A., Zhang, H., Clark, K. J., Ekker, S. C., & Wen, X.-Y. (2017). Disruption of pdgfra alters endocardial and myocardial fusion during zebrafish cardiac assembly. Biology Open, bio.021212. 10.1242/bio.021212

24. Fredriksson, L., Li, H., & Eriksson, U. (2004). The PDGF family: Four gene products form five dimeric isoforms. Cytokine & Growth Factor Reviews, 15(4), 197–204. 10.1016/j.cytogfr.2004.03.007

25. Grotendorst, G. R., Seppä, H. E., Kleinman, H. K., & Martin, G. R. (1981). Attachment of smooth muscle cells to collagen and their migration toward platelet-derived growth factor. Proceedings of the National Academy of Sciences, 78(6), 3669–3672. 10.1073/pnas.78.6.3669

26. Guérit, E., Arts, F., Dachy, G., Boulouadnine, B., & Demoulin, J.-B. (2021). PDGF receptor mutations in human diseases. Cellular and Molecular Life Sciences, 78(8), 3867–3881. 10.1007/s00018-020-03753-y

27. Hansson, M. D., Rzeznicka, K., Rosenbäck, M., Hansson, M., & Sirijovski, N. (2008). PCR- mediated deletion of plasmid DNA. Analytical Biochemistry, 375(2), 373–375. 10.1016/j.ab.2007.12.005

28. Horikawa, S., Ishii, Y., Hamashima, T., Yamamoto, S., Mori, H., Fujimori, T., Shen, J., Inoue, R., Nishizono, H., Itoh, H., Majima, M., Abraham, D., Miyawaki, T., & Sasahara, M. (2015). PDGFRα plays a crucial role in connective tissue remodeling. Scientific Reports, 5(1), 17948. 10.1038/srep17948

29. Johnson, R. L., Laufer, E., Riddle, R. D., & Tabin, C. (1994). Ectopic expression of Sonic hedgehog alters dorsal-ventral patterning of somites. Cell, 79(7), 1165–1173. 10.1016/0092-8674(94)90008-6

30. Karlstrom, R. O., Tyurina, O. V., Kawakami, A., Nishioka, N., Talbot, W. S., Sasaki, H., & Schier, A. F. (2003). Genetic analysis of zebrafish gli1 and gli2 reveals divergent requirements for gli genes in vertebrate development. Development, 130(8), 1549– 1564. 10.1242/dev.00364

31. Kawakami, M., Umeda, M., Nakagata, N., Takeo, T., & Yamamura, K. (2011). Novel migrating mouse neural crest cell assay system utilizing P0-Cre/EGFP fluorescent time-lapse imaging. BMC Developmental Biology, 11(1), 68. 10.1186/1471-213X-11-68

32. Kettleborough, R. N. W., Busch-Nentwich, E. M., Harvey, S. A., Dooley, C. M., De Bruijn, E., Van Eeden, F., Sealy, I., White, R. J., Herd, C., Nijman, I. J., Fényes, F., Mehroke, S., Scahill, C., Gibbons, R., Wali, N., Carruthers, S., Hall, A., Yen, J., Cuppen, E., & Stemple, D. L. (2013). A systematic genome-wide analysis of zebrafish protein-coding gene function. Nature, 496(7446), 494–497. 10.1038/nature11992

33. Krishnamurthy, V. V., Khamo, J. S., Cho, E., Schornak, C., & Zhang, K. (2015). Polymerase chain reaction-based gene removal from plasmids. Data in Brief, 4, 75–82. 10.1016/j.dib.2015.04.024

34. Loh, K. M., Chen, A., Koh, P. W., Deng, T. Z., Sinha, R., Tsai, J. M., Barkal, A. A., Shen, K. Y., Jain, R., Morganti, R. M., Shyh-Chang, N., Fernhoff, N. B., George, B. M., Wernig, G., Salomon, R. E. A., Chen, Z., Vogel, H., Epstein, J. A., Kundaje, A., … Weissman, I. L. (2016). Mapping the Pairwise Choices Leading from Pluripotency to Human Bone, Heart, and Other Mesoderm Cell Types. Cell, 166(2), 451–467. 10.1016/j.cell.2016.06.011

35. Ma, R. C., Jacobs, C. T., Sharma, P., Kocha, K. M., & Huang, P. (2018). Stereotypic generation of axial tenocytes from bipartite sclerotome domains in zebrafish. PLOS Genetics, 14(11), e1007775. 10.1371/journal.pgen.1007775

36. Ma, R. C., Kocha, K. M., Méndez-Olivos, E. E., Ruel, T. D., & Huang, P. (2023). Origin and diversification of fibroblasts from the sclerotome in zebrafish. Developmental Biology, 498, 35–48. 10.1016/j.ydbio.2023.03.004

37. Mao, A., Li, Z., Ning, G., Zhou, Z., Wei, C., Li, J., He, X., & Wang, Q. (2023). Sclerotome- derived PDGF signaling functions as a niche cue responsible for primitive erythropoiesis. Development, 150(22), dev201807. 10.1242/dev.201807

38. Mao, A., Zhang, M., Liu, J., Cao, Y., & Wang, Q. (2019). PDGF signaling from pharyngeal pouches promotes arch artery morphogenesis. Journal of Genetics and Genomics, 46(12), 551–559. 10.1016/j.jgg.2019.11.004

39. McDonald, J. A., Pinheiro, E. M., & Montell, D. J. (2003). PVF1, a PDGF/VEGF homolog, is sufficient to guide border cells and interacts genetically with Taiman. Development, 130(15), 3469–3478. 10.1242/dev.00574

40. Montero, J.-A., Kilian, B., Chan, J., Bayliss, P. E., & Heisenberg, C.-P. (2003). Phosphoinositide 3-Kinase Is Required for Process Outgrowth and Cell Polarization of Gastrulating Mesendodermal Cells. Current Biology, 13(15), 1279–1289. 10.1016/S0960-9822(03)00505-0

41. Mueller, A. A., Van Velthoven, C. T., Fukumoto, K. D., Cheung, T. H., & Rando, T. A. (2016). Intronic polyadenylation of PDGFRα in resident stem cells attenuates muscle fibrosis. Nature, 540(7632), 276–279. 10.1038/nature20160

42. Muhl, L., Genové, G., Leptidis, S., Liu, J., He, L., Mocci, G., Sun, Y., Gustafsson, S., Buyandelger, B., Chivukula, I. V., Segerstolpe, Å., Raschperger, E., Hansson, E. M., Björkegren, J. L. M., Peng, X.-R., Vanlandewijck, M., Lendahl, U., & Betsholtz, C. (2020). Single-cell analysis uncovers fibroblast heterogeneity and criteria for fibroblast and mural cell identification and discrimination. Nature Communications, 11(1), 3953. 10.1038/s41467-020-17740-1

43. Murayama, E., Sarris, M., Redd, M., Le Guyader, D., Vivier, C., Horsley, W., Trede, N., & Herbomel, P. (2015). NACA deficiency reveals the crucial role of somite-derived stromal cells in haematopoietic niche formation. Nature Communications, 6(1), 8375. 10.1038/ncomms9375

44. Murayama, E., Vivier, C., Schmidt, A., & Herbomel, P. (2023). Alcam-a and Pdgfr-α are essential for the development of sclerotome-derived stromal cells that support hematopoiesis. Nature Communications, 14(1), 1171. 10.1038/s41467-023-36612-y

45. Ono, Y., Yu, W., Jackson, H. E., Parkin, C. A., & Ingham, P. W. (2015). Adaxial cell migration in the zebrafish embryo is an active cell autonomous property that requires the Prdm1a transcription factor. Differentiation, 89(3–4), 77–86. 10.1016/j.diff.2015.03.002

46. Orr-Urtreger, A., & Lonai, P. (1992). Platelet-derived growth factor-A and its receptor are expressed in separate, but adjacent cell layers of the mouse embryo. Development, 115(4), 1045–1058. 10.1242/dev.115.4.1045

47. Pickett, E. A., Olsen, G. S., & Tallquist, M. D. (2008). Disruption of PDGFRα-initiated PI3K activation and migration of somite derivatives leads to spina bifida. Development, 135(3), 589–598. 10.1242/dev.013763

48. Plikus, M. V., Wang, X., Sinha, S., Forte, E., Thompson, S. M., Herzog, E. L., Driskell, R. R., Rosenthal, N., Biernaskie, J., & Horsley, V. (2021). Fibroblasts: Origins, definitions, and functions in health and disease. Cell, 184(15), 3852–3872. 10.1016/j.cell.2021.06.024

49. Rajan, A. M., Ma, R. C., Kocha, K. M., Zhang, D. J., & Huang, P. (2020). Dual function of perivascular fibroblasts in vascular stabilization in zebrafish. PLOS Genetics, 16(10), e1008800. 10.1371/journal.pgen.1008800

50. Rajan, A. M., Rosin, N. L., Labit, E., Biernaskie, J., Liao, S., & Huang, P. (2023). Single-cell analysis reveals distinct fibroblast plasticity during tenocyte regeneration in zebrafish. Science Advances, 9(46), eadi5771. 10.1126/sciadv.adi577

51. Scaal, M. (2016). Early development of the vertebral column. Seminars in Cell & Developmental Biology, 49, 83–91. 10.1016/j.semcdb.2015.11.003

52. Schindelin, J., Arganda-Carreras, I., Frise, E., Kaynig, V., Longair, M., Pietzsch, T., Preibisch, S., Rueden, C., Saalfeld, S., Schmid, B., Tinevez, J.-Y., White, D. J., Hartenstein, V., Eliceiri, K., Tomancak, P., & Cardona, A. (2012). Fiji: An open-source platform for biological-image analysis. Nature Methods, 9(7), 676–682. 10.1038/nmeth.2019

53. Seppa, H., Grotendorst, G., Seppa, S., Schiffmann, E., & Martin, G. R. (1982). Platelet- derived Growth Factor Is Chemotactic for Fibroblasts. The Journal of Cell Biology, 92, 584–588. 10.1083/jcb.92.2.584

54. Soriano, P. (1997). The PDGFα receptor is required for neural crest cell development and for normal patterning of the somites. Development, 124(14), 2691–2700. 10.1242/dev.124.14.2691

55. Stickney, H. L., Barresi, M. J. F., & Devoto, S. H. (2000). Somite Development in Zebrafish. Developmental Dynamics, 219(3), 287–303.

56. Tallquist, M. D. (2020). Cardiac Fibroblast Diversity. Annual Review of Physiology, 82(1), 63–78. 10.1146/annurev-physiol-021119-034527

57. Tallquist, M. D., & Soriano, P. (2003). Cell autonomous requirement for PDGFRα in populations of cranial and cardiac neural crest cells. Development, 130(3), 507–518. 10.1242/dev.00241

58. Tallquist, M. D., Weismann, K. E., Hellström, M., & Soriano, P. (2000). Early myotome specification regulates PDGFA expression and axial skeleton development. Development, 127(23), 5059–5070. 10.1242/dev.127.23.5059

59. Thisse, C., & Thisse, B. (2008). High-resolution in situ hybridization to whole-mount zebrafish embryos. Nature Protocols, 3(1), 59–69. 10.1038/nprot.2007.514

60. Yamaguchi, N., Colak-Champollion, T., & Knaut, H. (2019). zGrad is a nanobody-based degron system that inactivates proteins in zebrafish. eLife, 8, e43125. 10.7554/eLife.43125

61. Younesi, F. S., Miller, A. E., Barker, T. H., Rossi, F. M. V., & Hinz, B. (2024). Fibroblast and myofibroblast activation in normal tissue repair and fibrosis. Nature Reviews Molecular Cell Biology, 25, 617–638. 10.1038/s41580-024-00716-0

62. Zhang, Y., Werling, U., & Edelmann, W. (2012). SLiCE: A novel bacterial cell extract-based DNA cloning method. Nucleic Acids Research, 40(8), e55–e55. 10.1093/nar/gkr1288

